# How long is the brain perfusable after global ischemia? A systematic review

**DOI:** 10.64898/2026.07.10.737845

**Authors:** Jeremy Kalfus, Devin Ward, Borys Wróbel, Andrew T. McKenzie

## Abstract

**Background:** Global cerebral ischemia initiates a cascade of pathophysiological changes that progressively impair subsequent perfusion of brain tissue. Some authors have proposed that adequate cerebral perfusion becomes impossible after approximately 10-30 minutes of global ischemia. However, the extant evidence base for this threshold and its variation across studies have not yet been systematically examined.

**Objective:** To synthesize the literature on post-ischemic cerebral perfusion success as a function of ischemia duration.

**Methods:** We searched PubMed (February 11, 2026) for studies of global cerebral ischemia in animal or human models that reported quantitative or categorical measures of perfusion quality. Eligible studies included those assessing perfusion via restoration of blood flow, via external perfusion of non-blood solutions, and/or via tracer injection following reperfusion. Studies of focal ischemia were excluded. Data extracted included species, ischemia duration, temperature during ischemia, ischemia model, perfusate type, and perfusion quality assessment method. The perfusion quality outcome was operationalized as either the average percentage of brain tissue perfused or the percentage of brains in a group that were adequately perfused. Study quality was assessed using a custom domain-specific checklist. The review protocol was preregistered on the Open Science Framework (https://osf.io/2qm3w).

**Results:** We included 60 studies with 192 study arms reporting on the perfusion of the brains of rabbits, rats, pigs, dogs, cats, and humans. Studies differed in the model of ischemia, the perfusate, the perfusion parameters, the quality assessment methods, and other factors. Longer ischemia was associated with lower perfusion quality, but the relationship appeared to be a gradual decline rather than a sharp threshold. Additionally, the variability across studies was large, and some studies have found that at least partial perfusion is possible after longer periods. Within-study dose-response curves were more consistent than the pooled cross-study pattern.

**Conclusions:** How long the brain remains perfusable after circulatory arrest has not yet been definitively established. On average, perfusion quality clearly declines rapidly as the duration of global cerebral ischemia increases. However, some studies, often using interventions such as hypothermia or vasopressors, have reported at least partial perfusion of the brain even after 30 or 60 minutes of ischemia. Moreover, at least partial perfusion has been reported in human brain banking studies after postmortem intervals of several hours or days in some donors. Limitations of this review include substantial heterogeneity in study methods and outcome measures, which precluded formal meta-analysis. Future research may benefit from more thorough and precise measures of perfusion quality.

## Introduction

Brain banking is a critical foundation for neuroscience and neuropathology research. One important method sometimes used in brain banking is the use of machine perfusion to deliver preservative chemicals, such as aldehyde fixatives, through the brain’s vasculature system. This enables more rapid and uniform tissue preservation compared to immersion-based approaches [1,2]. With recent growth in the field of connectomics, which aims to profile the synaptic connectivity of larger areas of human brain tissue in the coming years, there is a critical need to improve our ability to perfuse postmortem brains [3]. Machine perfusion is also relevant to the emerging field of structural brain preservation, which is based upon the long-term preservation of the brain with the goal of maintaining the structural information that encodes memories and personal identity [4,5]. However, the effectiveness of perfusion-based brain preservation depends critically on the ability to deliver solutions through the microvasculature of the brain, which is progressively compromised after circulatory arrest. As the ischemic time increases after circulatory arrest, a constellation of pathological changes will progressively impair the vascular flow and thereby reduce the quality of perfusion [6–8].

The question of how rapidly cerebral perfusability is lost after circulatory arrest is also relevant to research in other fields in addition to brain banking. For example, in resuscitation science, understanding whether adequate cerebral reperfusion can be achieved after varying intervals of cardiac arrest is relevant to predicting neurological recovery [9]. In particular, the no-reflow phenomenon has been recognized as a contributor to poor neurological outcomes after cardiac arrest [10,11]. In the stroke and thrombectomy literature, data suggests that a substantial minority of patients with successful macrovascular recanalization can still have microvascular hypoperfusion [12]. Finally, in organ transplantation, particularly in donation after circulatory death protocols, understanding the window of perfusability informs organ viability decisions [13]; some mechanisms defining this window are expected to be common for different organs, even if the brain perfusion has unique features, e.g., because of the presence of the blood-brain barrier and because the brain volume is constrained by the skull.

Within the structural brain preservation literature, several specific perfusability thresholds have been proposed. For example, Michael Darwin has written that widespread microcirculatory failure occurs approximately 15 minutes after the onset of normothermic ischemia and is a “relatively sharp threshold beyond which effective cryoprotective perfusion of the brain becomes improbable” [14]. He also notes that once normothermic ischemia has exceeded 30 minutes, perfusing the cerebral vasculature for washout is essentially unachievable, unless tightly controlled reperfusion methods are used [14]. As another example, one study in rats found that blood washout with the organ preservation solution MHP-2 substantially ameliorated perfusion impairment following 30 minutes of warm ischemia compared to leaving blood in the vasculature [15]. A benefit from perfusion-based washout was observed across warm ischemic intervals up to 50 minutes, but at 60 minutes and beyond, complete freezing of the brain occurred regardless of whether washout was performed, indicative of inadequate cryoprotectant perfusion [15]. One source reported that in rat experiments using aldehyde-stabilized cryopreservation, perfusion initiated within 12 minutes of death yielded “textbook-quality” preservation, while longer delays resulted in segments of blood vessels failing to perfuse [16]. Finally, one study reported a perfusability window of approximately 14 minutes in adult pigs undergoing aldehyde-stabilized cryopreservation [17]. This was based on successful ultrastructural preservation in one pig after 13.8 minutes of ischemia and the failure of preservation in two pigs after 18.3 and 22.8 minutes of ischemia, which the authors attributed to having exceeded the perfusability window [17].

Studies across several decades have measured perfusion quality after varying durations of ischemia, and a systematic review of this evidence base could help to ground the discussion of perfusability windows in a summary of the existing data. In this review, we set out to review this literature by gathering evidence from animal and human studies which characterize how perfusion quality changes as a function of the duration of global cerebral ischemia. Our goal is to provide an evidence base that may help to inform decisions in brain banking and brain preservation, while also potentially being useful to researchers and clinicians in other fields.

## Review Methods

### Study design

The protocol for the review was pre-registered at Open Science Framework (https://osf.io/2qm3w/files/7rg6j). We report our adherence to the PRISMA 2020 systematic review reporting standards [18] (**Data S1**).

### Search strategy

We searched PubMed on February 11, 2026 using the following query:

> “brain” AND (“perfusion” OR “reperfusion”) AND (“ischemia” OR “postmortem” OR “brain banking” OR “autopsy”) AND (“carbon black” OR “evans blue” OR “india ink” OR “trypan blue” OR “fluorescein” OR “horseradish peroxidase” OR “microspheres” OR “FITC-dextran” OR “no reflow” OR “vascular patency” OR “microvascular obstruction”)

This yielded 894 abstracts, which were uploaded to SysRev for screening. We supplemented the PubMed search with *ad hoc* searches and forward/backward citation tracing, particularly from foundational studies (e.g. [10,11]).

### Eligibility Criteria

#### Inclusion criteria

Studies were included if they report empirical data on brain perfusion quality, or on some metric that could be used to infer brain perfusion quality, following a defined period of global cerebral ischemia. Eligible studies include those assessing perfusion via restoration of blood flow (e.g., after cardiac resuscitation or release of vascular occlusion), via external perfusion of non-blood solutions (e.g., saline or tracer suspensions), and/or via tracer injection following reperfusion. Studies on humans or non-human animals of any age were included. Studies were grouped by perfusate type (e.g., blood vs. non-blood) and species for subgroup analyses.

We required the ischemic insult to be both complete (no residual cerebral blood flow) and global (affecting the whole brain). We therefore excluded focal and regional models such as middle cerebral artery occlusion, models leaving potential collateral flow such as four-vessel occlusion in the rat [19], or models leaving vertebrobasilar flow such as bilateral carotid occlusion alone. More generally, we excluded any model in which the paper reported or accepted residual cerebral blood flow during the ischemic period. Where a study included a retrograde cerebral perfusion arm, we extracted the data only from the standard circulatory arrest group.

#### Exclusion criteria

Focal/regional ischemia models (e.g., middle cerebral artery occlusion); reviews or editorials without original data; case reports; articles not available in English.

#### Screening and Selection

Each title and abstract was screened by at least one reviewer to identify potentially eligible studies (**Data S2**). Next, at least one reviewer reviewed the full-text of each identified abstract for potential inclusion in the data extraction process (**Data S3**). Disagreements on the inclusion of any articles reviewed by multiple people were resolved by discussion. A PRISMA flow diagram was used to document the study selection process.

### Data extraction

We treated each distinct combination of experimental group and assessment as a separate observation or study arm, so that a single study could have multiple study arms. Treated and untreated groups, groups subjected to different ischemia durations, and the same cohort assessed by different methods each constituted a separate study arm. From each study arm we extracted 27 data variables. Data variables included ischemia duration, species, sample size, temperature during ischemia, ischemia model, perfusate type, perfusion pressure, assessment method, perfusion outcome, variability, percent of animals adequately perfused, regional perfusion breakdown, brain regions assessed, and time from ischemia onset to assessment. For each study arm we also extracted interpretive variables, including the estimated percentage of the brain that was spatially perfused and a perfusion quality category.

We derived two measures of perfusion quality. First, we derived an estimate of the spatial perfusion completeness. This refers to the percentage of the brain that received any flow, as distinct from flow magnitude. For studies that report the non-perfused area, we computed 100% minus that area; for ordinal filling grades, we used the grade midpoints averaged across animals; and for studies that report the fraction of microvasculature reperfused, we used the reported value. Studies that did not report whole brain spatial coverage, such as those measuring perfusion with single-site flow probes or large-vessel angiography, were not converted. Second, we derived a measure of the proportion of animals in a group whose perfusion was adequate, using each study’s own adequacy criterion where it was defined, and otherwise a default threshold of at least 90% spatial perfusion per animal, where the study did not define such a criterion.

Notably, spatial perfusion completeness can only be assessed at the spatial resolution that each method allows. Some methods (such as carbon black) can detect the degree of spatial perfusion completeness at the level of the microvasculature, while others (such as microsphere cerebral blood flow) can only detect whether each sampled region received at least some degree of flow. As a result, a given degree of spatial completeness of perfusion is stronger evidence of complete perfusion when measured by a method capable of resolving the microvasculature, because such a method can detect gaps within a larger region that received at least some flow.

For studies reporting several time points after the initiation of perfusion, we extracted the earliest stabilized measurement, because we are primarily interested in the initial post-ischemic state before secondary processes resolve or worsen it. In the cases where the spatial perfusion pattern changed qualitatively over time, we also recorded these later qualitative changes for descriptive purposes. In cardiac arrest models with cardiopulmonary resuscitation, we counted only the pre-resuscitation no-flow interval as the ischemia duration, since chest compressions are expected to produce partial circulation.

To assist with extraction, we developed an LLM-based extraction prompt through a series of iterative tests and refinements. First, the data were extracted twice using separate LLM instances (Claude Opus 4.6, Anthropic; sessions with extended thinking enabled), each receiving the same structured prompt and the full-text PDF of the study. Next, the two resulting spreadsheets were compared by a third LLM session, which produced a discrepancy report flagging disagreements. Based on any discrepancies noted, the prompt was edited, in an iterative process, with the aim to remove such discrepancies as much as possible. Convergence to a minimal number of discrepancies signified that a final prompt was obtained. Then, each included paper underwent data extraction using this final prompt (available in **Data S4**).

For each study, we assessed study quality using a custom set of indicators. To the best of our knowledge, there are no validated risk-of-bias tools for studies of this type, so we developed a domain-specific checklist. For each study, recorded whether: (1) the ischemia duration was clearly defined and experimentally controlled, (2) the sample size was explicitly reported for each group, (3) the temperature during ischemia was reported, (4) the perfusion outcome assessment was described in sufficient detail to be reproducible, (5) any blinding was used in outcome assessment, and (6) the inclusion or exclusion criteria for subjects were stated. Each indicator was coded as yes, no, or unclear. Study quality was summarized descriptively across the included studies.

A second reviewer performed a manual check of the data extracted by the AI in a randomly selected subset of studies (12/60, 20%, using random.org), covering both the data extraction and study quality assessments. For data extraction, we had 87.4% agreement (1,682/1,924 data points), 10.0% non-substantive disagreements (193/1,924 data points; e.g., incorrect formatting but correct data), and 2.5% substantive disagreements (49/1,924 data points; **Data S5**). For study quality, we had 99.5% agreement (203/204 data points), 0.0% non-substantive disagreement (0/204 data points), and 0.5% (1/204 data points) substantive disagreement (**Data S5**). For both the data extraction and study quality assessment, this level of agreement exceeded our pre-specified threshold for at least 80% agreement (i.e., a substantive disagreement rate of 20% or less), indicating an acceptable degree of accuracy for data extraction, consistent with thresholds commonly used in other systematic reviews [20].

### Data Synthesis

Our included studies, study quality assessments, and extracted data are available (**Data S6**). For all statistical analysis, we used R (v. 4.5.3). We synthesized the data qualitatively and graphically. We plotted ischemia duration (x-axis) against perfusion quality (y-axis), with the points being annotated by different potential moderating variables such as perfusate type (i.e., blood vs non-blood). These plots include all available data points, regardless of whether the study arm used an intervention to improve perfusion quality. We also qualitatively examined potential moderating factors. No formal statistical assessment of reporting bias (e.g., funnel plots) was performed, as no meta-analysis was conducted.

We assessed our confidence in each major finding using a structured narrative approach, based on qualitative factors including (a) the number of supporting studies, (b) the consistency of findings across studies, (c) whether studies had direct perfusion measurement or more indirect outcomes, (d) the study quality, and (e) the consistency across species. Confidence was characterized as high, moderate, low, or very low for each major finding, and presented in a summary table alongside the main conclusions.

### Differences between the protocols and the review

Relative to the pre-registered protocols for this review (available at https://osf.io/2qm3w/files/osfstorage), we made several changes. We registered three protocol versions on OSF, and the changes below describe how the final review substantively departs from our original plans described in one or more of these pre-registered protocols.

First, we changed the review type. The first registered version described a realist synthesis. We switched to a systematic review before completing the abstract screening process, because we decided this was a better fit for the more focused question we were considering here.

Second, we broadened the eligibility criteria. The first version restricted inclusion to studies using non-blood perfusates or modified blood reperfusion following a washout step, and it excluded studies using unmodified whole blood reperfusion. We removed that restriction and included studies that perfused with blood after global cerebral ischemia, because the non-blood perfusate literature turned out to be smaller than we expected once we started screening, and the blood reperfusion studies turned out to be relevant to the same question.

Third, we changed how we validated the data extraction. The protocols stated that all extracted entries would be manually reviewed and verified against the data in the source paper. Instead, one reviewer manually reviewed a randomly selected subset of studies, which we decided was sufficient to estimate extraction accuracy. We also did not complete the planned comparison of AI-assisted extraction between two independent reviewers. This is because we decided that the iterative refinement of the extraction prompt we performed, combined with the manual verification of the extracted data in a random subset of studies, provided more meaningful validation than comparing two independent AI-assisted extraction results.

Fourth, we simplified the planned analyses of study quality. After assessing study quality using a domain-specific checklist, we found that quality scores showed limited variation across studies and were also confounded with other methodological characteristics. We therefore concluded that sensitivity analyses based on study quality would be unlikely to yield meaningful insights and instead summarized these assessments descriptively.

## Results and Discussion

### Study selection

We identified 60 full text articles that had a degree of global cerebral ischemia followed by a brain perfusion assessment (**Figure 1**). From these 60 studies, we extracted 192 observations with distinct experimental conditions, which we designated as study arms. In terms of species, rabbits were the most common (66 study arms), followed by rats (36), pigs (31), dogs (31), cats (25), and humans (3). Ischemia durations ranged from 1 to 60 minutes (median 15 minutes), excluding one outlier of human donor cases with an average postmortem interval of around 37 hours, which is analyzed separately [2]. Of the 192 arms, 121 (63%) used blood as the perfusate at the time of assessment, 67 (35%) used a non-blood solution, and 4 (2%) used a mixture of the two, either residual blood incompletely displaced by a non-blood perfusate or a deliberate mix of blood and a synthetic hemoglobin-based solution.

**Figure 1.**
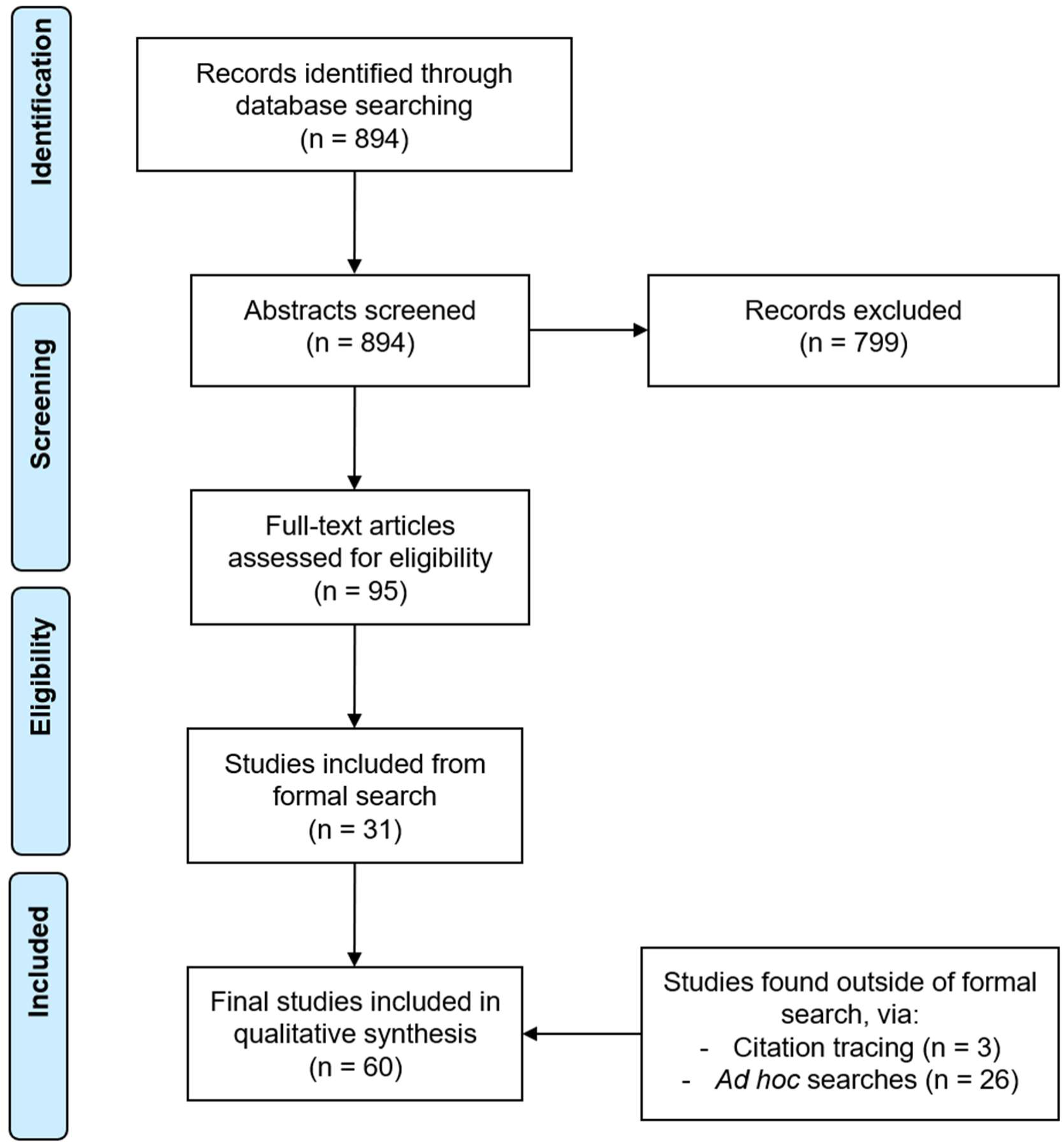
Study selection diagram. In the included studies, perfusion quality was assessed using a range of methods. The most common assessment method was carbon black perfusion, with India ink being a common formulation used. In this method, a particulate tracer (carbon particles up to a few microns in size) is added to the perfusate, and perfused or non-perfused regions are identified by the extent to which the tracer fills the cerebral vasculature, scored either on the brain surface or in coronal sections (n = 68 study arms).

Quantitative cerebral blood flow (CBF) was measured in a substantial number of arms, through a variety of methods. First, by radiolabeled or fluorescent microspheres, which become trapped in the microvasculature in proportion to regional flow (n = 28 arms). Second, by antipyrine or iodoantipyrine autoradiography, which maps regional CBF based on the tissue distribution of a diffusible radiolabeled tracer (n = 14 arms). Third, by xenon-enhanced CT (Xe-CT), which images the wash-in of xenon to produce voxel-level CBF maps (n = 14 arms). Finally, via 133-Xe clearance, which derives CBF from the washout rate of radioactive xenon measured by external detectors (n = 7 arms).

Filling of the microvasculature was visualized directly by fluorescein isothiocyanate (FITC)-albumin fluorescence microscopy, in which a fluorescent plasma tracer labels perfused vessels (n = 12 arms), and by intravital imaging of functional capillary density, including sidestream dark-field microscopy, which quantifies the length of perfused capillaries per a given tissue area (n = 8 arms). Surface blood flow was estimated optically by laser Doppler or laser speckle flowmetry from the motion of red cells (n = 10 arms). Several other methods, including angiography (n = 5 arms), accounted for the remainder.

### Perfusion quality changes with ischemia time

We operationalized perfusion quality from the heterogeneous outcomes reported across studies onto a common scale via two measures. First, we derived an estimate of the percentage of brain tissue that was spatially perfused, either for the single brain in a study arm or averaged across the brains in the arm. Second, we derived the proportion of brains in the arm that met a threshold for adequate perfusion. Depending on the data reported in the study, we were able to derive one, both, or neither of these metrics for each study or study arm.

We found that across studies, the extent of spatial perfusion across the brain declined with ischemia duration (**Figure 2**). Studies in which spatial perfusion was assessed used either carbon black or pure blood as perfusate. Interestingly, the decline was present predominantly in the arms using carbon black, and was accompanied by a wide degree of variability at any given duration. Carbon black filling fell from consistently near complete at very short durations to a broad spread by 15 minutes, with individual arms ranging from roughly 5% to 90% perfused. Study arms perfusing blood, on the other hand, had higher spatial perfusion completeness across the ischemia time points studied, with clusters of data points near 100% even after 30 to 60 minutes of ischemia, in some experimental paradigms [21–24].

**Figure 2.**
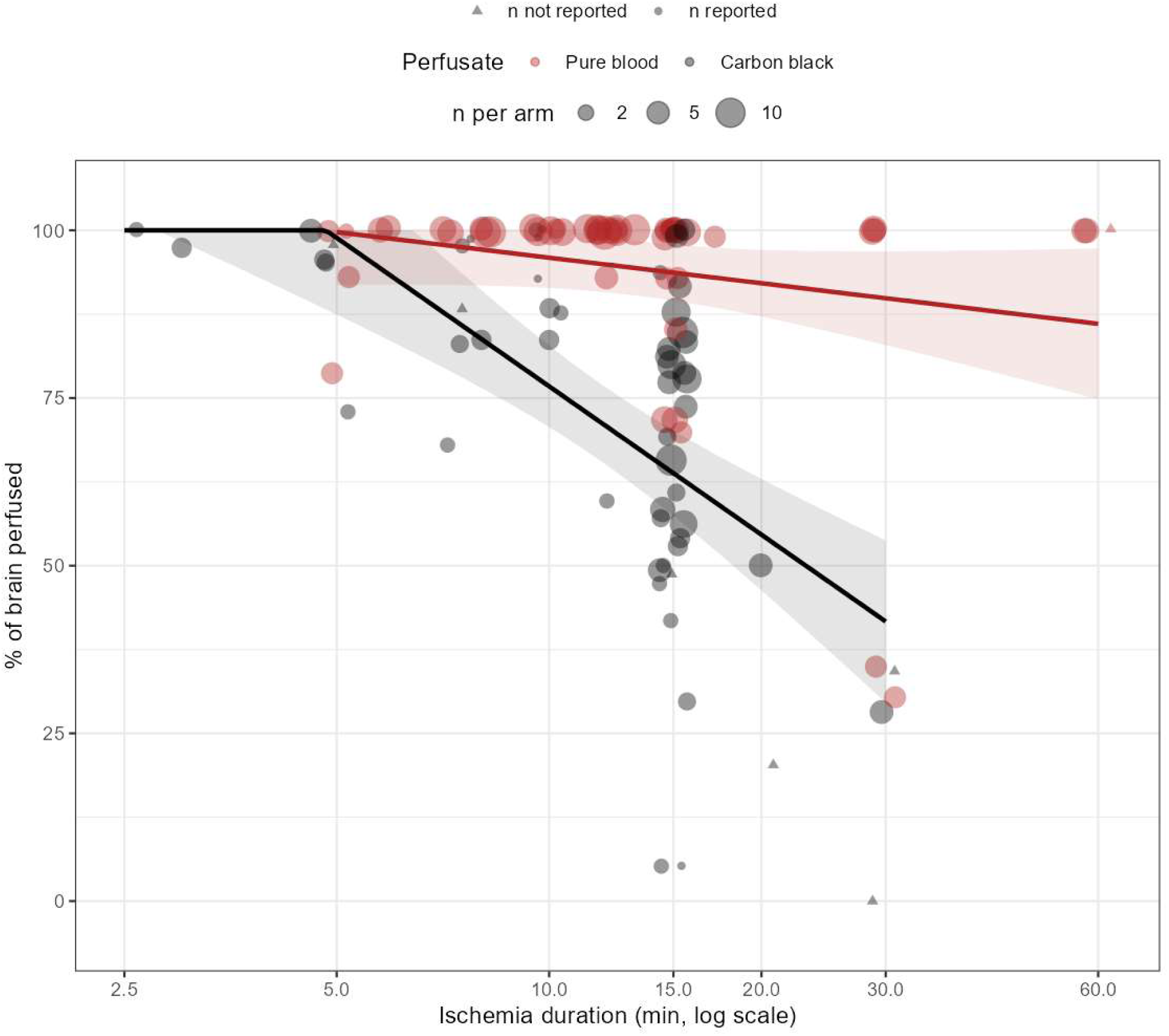
Spatial completeness of brain perfusion after different durations of global ischemia. Each point represents a study arm. The x-axis shows ischemia duration (log scale), and the y-axis shows the estimated percentage of the brain that was perfused. Point size is proportional to sample size, point color indicates perfusate type, and point shape indicates whether the sample size was reported. Points are shown with slight horizontal jitter and partial transparency to reduce overplotting. Solid lines show linear regression lines fit against log10-transformed ischemia duration for the pure blood and carbon black groups, with shaded bands indicating 95% confidence intervals. An interactive version of this figure is available here: https://andrewtmckenzie.com/Ischemic_perfusion_review/spatial-completeness-of-perfusion.html and in **Data S7**.

There are multiple potential explanations for the discrepancy between the carbon black and blood study arms. First, the perfusate type is confounded with how the perfusion was done. The carbon black arms are often open-circuit, at a fixed pressure, and with relatively low volume infusions (e.g. 50 mL of perfusate [23]). On the other hand, the blood arms are largely *in vivo* reperfusion studies in which the heart provides continuous circulation, and the assessment frequently occurred after a recirculation interval during which any initial perfusion deficits could partially resolve. Second, the quality assessment methods tend to differ. Carbon black is more sensitive to lower spatial completeness than the macroscopic assessment methods that were more commonly used in the blood perfusion arms. Third, properties of the carbon black perfusate may itself cause some of the defects. Multiple studies have reported deficits of cerebral perfusion after arterial infusion of solutions containing carbon black in control animals, even in the absence of prior ischemia [25,23]. This may be due to aggregates containing carbon black particles blocking blood vessels, which could be worse when the suspension is unfiltered [25]. Fourth, relatively more of the blood arms use hypothermia or other interventions such as vasopressors, which would be expected to improve perfusion quality [24,26]. As a result, we interpret this difference as related to the assessment method, experimental context, and potential problems with carbon black in particular, rather than indicating that there is something special about blood that makes it better for perfusion. Indeed, there is good reason to think that red blood cells would make perfusion more difficult, as they are relatively large. Moreover, other components in the blood may also contribute to thromboembolism or reperfusion injury during prolonged reperfusion.

As with the spatial completeness of perfusion, we found that the proportion of brains in a given arm that were adequately perfused decreased with increasing ischemic interval. Here also, there was a faster decline in the arms using carbon black (**Figure 3**). The decline for this frequency of adequate perfusion measure was more substantial than for the spatial completeness measure, with many data points at 0% beyond 10 minutes of ischemia. This is most likely due to the coding of the two measures rather than to a difference in the actual perfusion quality among the data points that each measure covers. Specifically, the spatial completeness measure captures partial perfusion across the brain as a continuous value, whereas the frequency of adequate perfusion measure applies a per-animal threshold that potentially scores a partially perfused brain as inadequate. This effectively thresholds a graded decline in the degree of spatial perfusion into a sharp drop in the proportion of brains classified as adequate.

**Figure 3.**
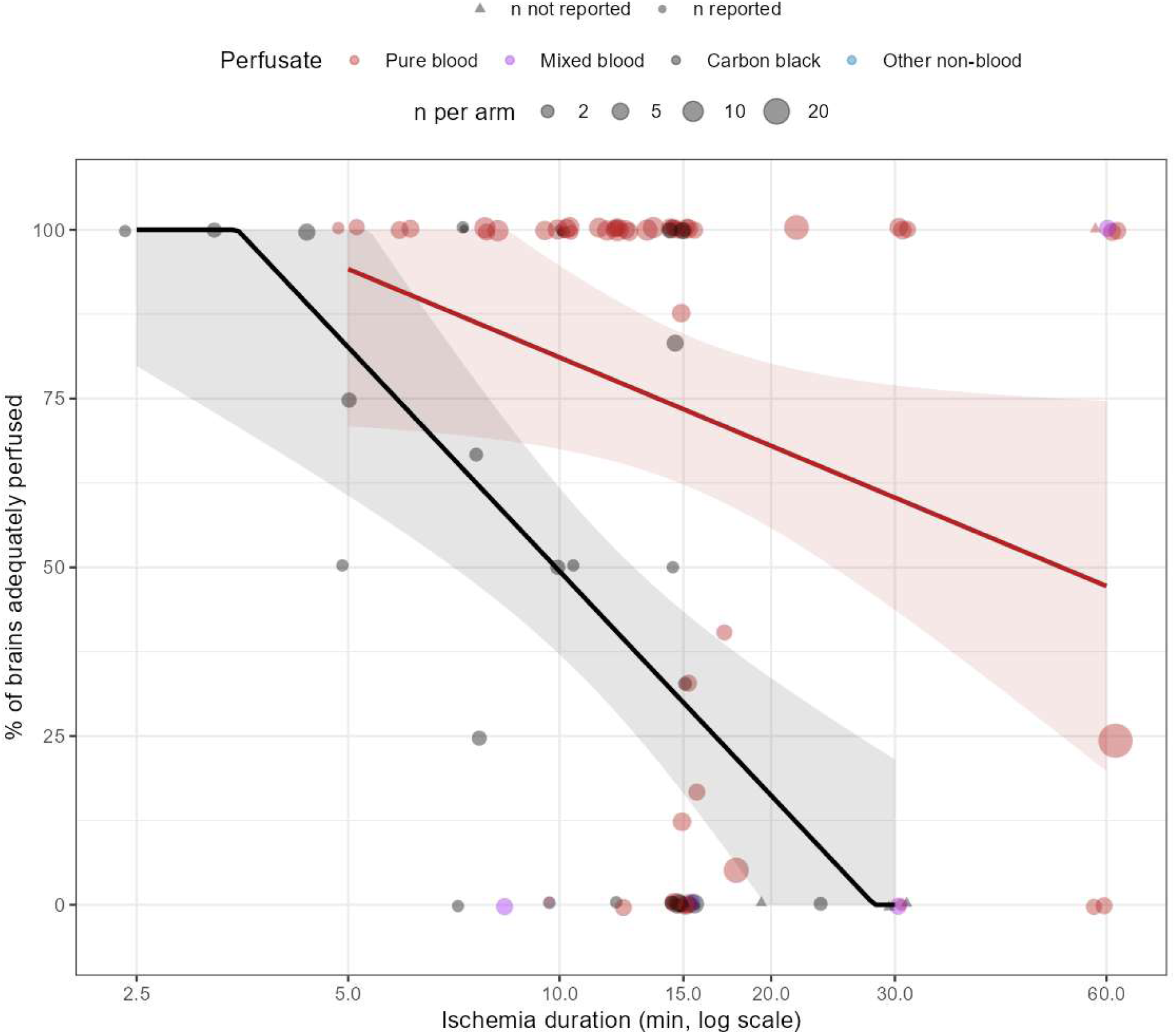
Frequency of adequate brain perfusion after different durations of global ischemia. Each point represents a study arm. The x-axis shows ischemia duration (log scale), and the y-axis shows the percentage of brains in each study arm that met the criterion for adequate perfusion. Point size is proportional to sample size, point color indicates perfusate type, and point shape indicates whether the sample size was reported. Points are shown with slight horizontal jitter and partial transparency to reduce overplotting. Solid lines show linear regression fits with ischemia duration modeled on the log10 scale for the pure blood and carbon black groups, with shaded bands indicating 95% confidence intervals. An interactive version of this figure is available here: : https://andrewtmckenzie.com/Ischemic_perfusion_review/frequency-of-adequate-perfusion.html and in **Data S8.**

### Within-study comparisons

Some of the included studies varied the ischemia duration while holding species, perfusate type, and assessment method constant. These within-study trends tended to show a more consistent decline in perfusion quality with longer ischemia than our approach of pooling data across studies. We describe a few examples of this here.

First, in the original study describing the no-reflow phenomenon, carbon black was perfused through the carotid arteries of rabbits after 2.5 to 15 minutes of ischemia [10]. The authors found that the nonperfused area rose with ischemia duration, reaching up to 95 percent of the brain in some animals after 15 minutes [10]. Second, in a similar rabbit model using carbon black perfused through the aorta, perfusion was complete below five minutes of ischemia, small perfusion deficits appeared by seven and a half minutes, roughly half the brain failed to perfuse by fifteen minutes, and the non-perfused area rose to about 60 percent by thirty minutes of ischemia [27]. Third, one study of rabbits, using carbon black perfused through the ascending aorta, framed the perfusion deficit in terms of the pressure needed to overcome it, and found that the minimum perfusion pressure required to fill the entire brain rose with the duration of ischemia [28]. They found that after only 5 minutes of ischemia, 95% of the brain reperfused even at a low pressure of only 22 mmHg, whereas the pressure needed to achieve complete reperfusion climbed to above roughly 50 mmHg after 10 minutes, above 70 mmHg after 15 minutes, and above about 103 mmHg after 30 minutes [28]. Fourth, in a cardiac arrest model, the cortical blood flow generated by external CPR in rabbits fell from about 28% of control after one minute of arrest to 9%, 6%, and 2% of control after three, five, and seven minutes, respectively, with no recordable flow after nine minutes [29]. Fifth, in normothermic cats whose circulating blood was labeled with FITC-albumin, forebrain no-reflow during CPR with epinephrine rose from about 21% of sectional area after five minutes of cardiac arrest to 42% and 70% after fifteen and thirty minutes, respectively [11].

Taken together, the contrast between these consistent within-study dose-response curves and the much noisier pooled relationship suggests that most of the cross-study variability at any given duration comes from methodological differences between studies rather than from biological variability in how rapidly the brain becomes unperfusable.

### Qualitative reperfusion outcomes

Some study arms reported perfusion outcomes that could not be converted into our percentage of brain perfused measure. For these arms, we instead classified the observations into different qualitative perfusion outcome categories during data extraction. Of the 81 such arms with ischemia duration of 60 minutes or less, we classified 34 of them as showing flow that returned adequately (early reactive hyperemia or no-reflow reported as absent), 16 as showing flow that returned but remained reduced (hypoperfusion without frank no-reflow), and 31 as showing flow that failed to return in some or all regions (no-reflow reported as present). Arms in which flow returned adequately and arms in which it failed to return had similar median ischemia durations (12.5 and 15 minutes, respectively), and consistent with this, there was no statistically significant difference in duration between these study arms (Wilcoxon rank-sum test, p = 0.87). The hypoperfusion arms clustered at shorter durations (median 7.5 minutes). We interpret these qualitative results as broadly consistent with the finding that the perfusion quality assessment method, along with other aspects of study methodology, is what primarily determines whether perfusion is scored as adequate or as no-reflow after a given duration of ischemia across the studies that we identified.

### Perfusion at extended postmortem intervals in human brain banking

The single human study included was a brain banking study of 77 whole-body donors in which the postmortem brains were perfused *in situ* with formalin, primarily through the internal carotid arteries [2]. Perfusion quality was assessed by gross examination, CT imaging, and histological grading, and was found to decline as the postmortem interval increased. However, at least partial perfusion was still observed in some donors at postmortem intervals up to several days, with several cases at intervals of 144 to 192 hours retaining moderate-to-high gross perfusion grades in at least some regions, well beyond the ischemia durations examined in the animal studies in our dataset. This finding is consistent with the idea that whether perfusion is scored as successful depends heavily on the assessment method and the quality threshold applied, rather than ischemia duration determining a binary perfusable or non-perfusable state. Although not included in our study due to their differing search terms, other human brain banking papers have also reported that human brains are at least partially perfusable up to 24 hours or more in at least some cases [1,30]. Notably, what these human brain banking studies demonstrate is vascular accessibility for fixative delivery, as opposed to reinstating physiological perfusion.

### Certain interventions can extend the perfusable window substantially

Some studies using pharmacological intervention or hypothermia reported good perfusability at ischemia durations where untreated brains typically fail. For example, in one study piglets that were cooled to 18°C before undergoing 60 minutes of deep hypothermic circulatory arrest showed measurable flow across all brain regions by microsphere CBF upon reperfusion [24]. In one study, giving hypertonic mannitol or hypertonic glucose before a 15-minute ischemic period in rabbits reduced carbon black filling defects, from 51% in untreated controls to 8% for mannitol and 1.5% for glucose [31]. Notably, the authors speculated that glucose’s stronger effect may have been partly due to its acting as a metabolic substrate rather than osmolarity alone. As another example of an intervention, one study tested the effect of acute pre-ischemic hemodilution down to a hematocrit of 21-32%, from a baseline of 40-48% [32]. They found that this eliminated the perfusion deficits occurring in non-cortical regions after 15 minutes of ischemia, compared with extensive defects in those regions in untreated control animals perfused at the same pressure.

### Partial resolution of perfusion impairment over time

Several included studies found that initial no-reflow can partially resolve during early recirculation. One study found that forebrain no-reflow in cats declined from 21% during CPR to 7% after 30 minutes of spontaneous recirculation following 5 minutes of cardiac arrest [11]. However, this resolution was highly dependent on the duration of initial ischemia, as after 15 or 30 minutes of cardiac arrest, no-reflow was found to not significantly decline after a period of spontaneous recirculation. Another study observed a similar pattern in rats. In this study, after 15 minutes of complete cerebral ischemia, the authors found that 5 minutes of recirculation revealed patchy perfusion defects consistent with no-reflow, but by 60 minutes of recirculation, these defects had resolved [33]. On the other hand, when the initial period of ischemia was extended to 30 minutes, perfusion defects persisted even after 90 minutes of recirculation.

One study measured how vascular obstruction resolves over the post-ischemic period [27]. After 15 minutes of complete ischemia in rabbits, when carbon black perfusion was done immediately, 44% of the cross-sectional area (averaged across six coronal sections) failed to fill. However, this fell to 21% after 7.5 minutes of restored normotensive flow, 13% after 15 minutes, and 5% after 2 hours. They claim that much of the early post-ischemic perfusion impairment is therefore dynamic and reversible once oxygenated blood is resupplied at normal pressure, though full resolution takes time.

Notably, the resolution of no-reflow does not necessarily indicate a return to normal perfusion. Instead, it could indicate a transition between two distinct phases of post-ischemic perfusion impairment. The initial no-reflow phenomenon is thought to be characterized by vascular obstruction — including intravascular rouleaux formation, endothelial cell swelling, and other factors — with the severity proportional to the ischemia duration [6]. This gives way to a delayed hypoperfusion phase, which is driven by arteriolar vasoconstriction and a loss of endothelial vasodilatory capacity [6]. As an example of this, in the rat study described above, regional CBF values at 60 minutes of recirculation ranged from 20-80% of control, indicating that the resolution of focal no-reflow can be accompanied by a widespread hypoperfusion [33].

For this review, we focused on the earliest available post-reperfusion measurement for each study because it provides the most unambiguous estimate of the brain’s initial perfusability following ischemia. Although longer periods of perfusion could potentially restore flow to additional regions as the vascular obstruction resolves, there is no clear point at which this process reaches completion, making it difficult to define a standardized endpoint. Moreover, in many applications, delayed perfusion may lead to reduced tissue quality, because autolysis will continue during the intervening period, and prolonged perfusion may itself introduce tissue alterations. For these reasons, we considered the earliest perfusion quality measurement to be the most informative endpoint for this review. However, we note that a future analysis of the same or similar data could instead focus on the temporal evolution of perfusion quality during prolonged perfusion.

### Regional differences

Several studies report that perfusion defects are distributed unevenly across the brain rather than uniformly. For example, one study found that obstruction was most severe in caudal and deep structures such as the cerebellum, brainstem, caudate, and thalamus, while the cerebral cortex was relatively spared [34]. Another study observed a similar pattern, with no-reflow found in the striatum, thalamus, hippocampus, and deeper cortical layers, although they also noted pronounced variation both between animals and between the two hemispheres of the same brain [33]. In our human brain banking study, perfusion was patchy in nearly all cases, both across and within vascular territories, with the posterior circulation generally perfusing less completely than the anterior circulation [2]. Similarly, in our postmortem pig brain perfusion study (not included in this systematic review because it was published after our search), we noted that cerebral white matter, especially in watershed areas, showed preservation deficits [17].

In addition to regional patterns, several authors noted an apparently random component, with given small areas failing to perfuse in one brain yet perfusing normally in another despite similar or longer durations of ischemia [23,34]. One possibility is anatomical variations in the vasculature. Another proposed mechanism attributes this patchiness to ischemia-induced loss of autoregulation acting on top of preexisting regional differences in how readily different areas reperfuse after ischemia [28]. The idea is that autoregulation normally equalizes flow across regions, damping out local differences in resistance. When it is absent – which will occur in the ischemic brain – the distribution of flow is instead controlled by differences in passive resistance. As a result, the regions in which the residual high-viscosity blood is cleared drop to low resistance and carry most of the incoming flow. On the other hand, areas where the blood is not cleared retain high vascular resistance and are underperfused, as the cleared regions “steal” flow from them. This mechanism has the potential to amplify small initial differences in perfusability across the brain into the patchy pattern that is observed, and it may be especially pronounced if low flow rates are used.

### Study quality

We scored each of the 60 included studies across six methodological dimensions. Reporting of ischemia duration, sample size, and outcome description were nearly universal (98%, 95%, and 100% of studies). The temperature during ischemia was reported less consistently, in 67% of studies. Blinding was reported in only 15% and inclusion/exclusion criteria in 42%. On this basis, quality scores ranged from 2 to 6 with a median of 4. Study quality ratings had a weak but statistically significant positive correlation with publication year (Spearman ρ = 0.28, p = 0.03), suggesting some improvement in reporting quality over the decades covered by our search. However, most studies, including more recent ones, still do not report blinding or clear inclusion/exclusion criteria for the animals studied, which raises the possibility of publication bias.

### Reporting bias

We did not formally assess reporting bias because the included studies were highly heterogeneous, preventing us from performing a meta analysis and using associated methods such as funnel plots. Nevertheless, we consider reporting bias to be a potential limitation of the available evidence base presented here. Most of the included studies were exploratory basic science experiments, for which formal reporting standards and prespecified analyses are uncommon. Consequently, unsuccessful or inconclusive findings are likely to be underrepresented in the published literature. For example, unsuccessful attempts to improve post-ischemic perfusion may have been less likely to be published than successful ones.

### Limitations

This review has several limitations that warrant consideration. First, we did not perform a formal meta-analysis. The heterogeneity across studies in species, ischemia model, perfusate, perfusion parameters, and assessment method was too large to support pooling into a single effect estimate, and the outcome measures were too inconsistent to combine quantitatively. As a result, our data synthesis is qualitative. The cross-study patterns we describe should therefore be read as descriptive rather than as actual effect size estimates. Second, our two perfusion quality measures required substantial derivation from heterogeneous data sources. Third, the studies we included have many confounds preventing us from cleanly isolating the effect of ischemia duration on perfusion quality. Fourth, the literature on deep hypothermic circulatory arrest was almost entirely outside of our search strategy. Only one study in our dataset used profound hypothermia, so our findings are predominantly relevant only to normothermic or mildly hypothermic ischemia and should not be extended to contexts where organisms are hypothermic prior to cardiac arrest, where the perfusable window is almost certainly much longer.

### Summary table

We assessed our confidence in the key findings of our review using a structured narrative approach. The findings and our confidence in them are summarized in **Table 1**.

**Table 1.**
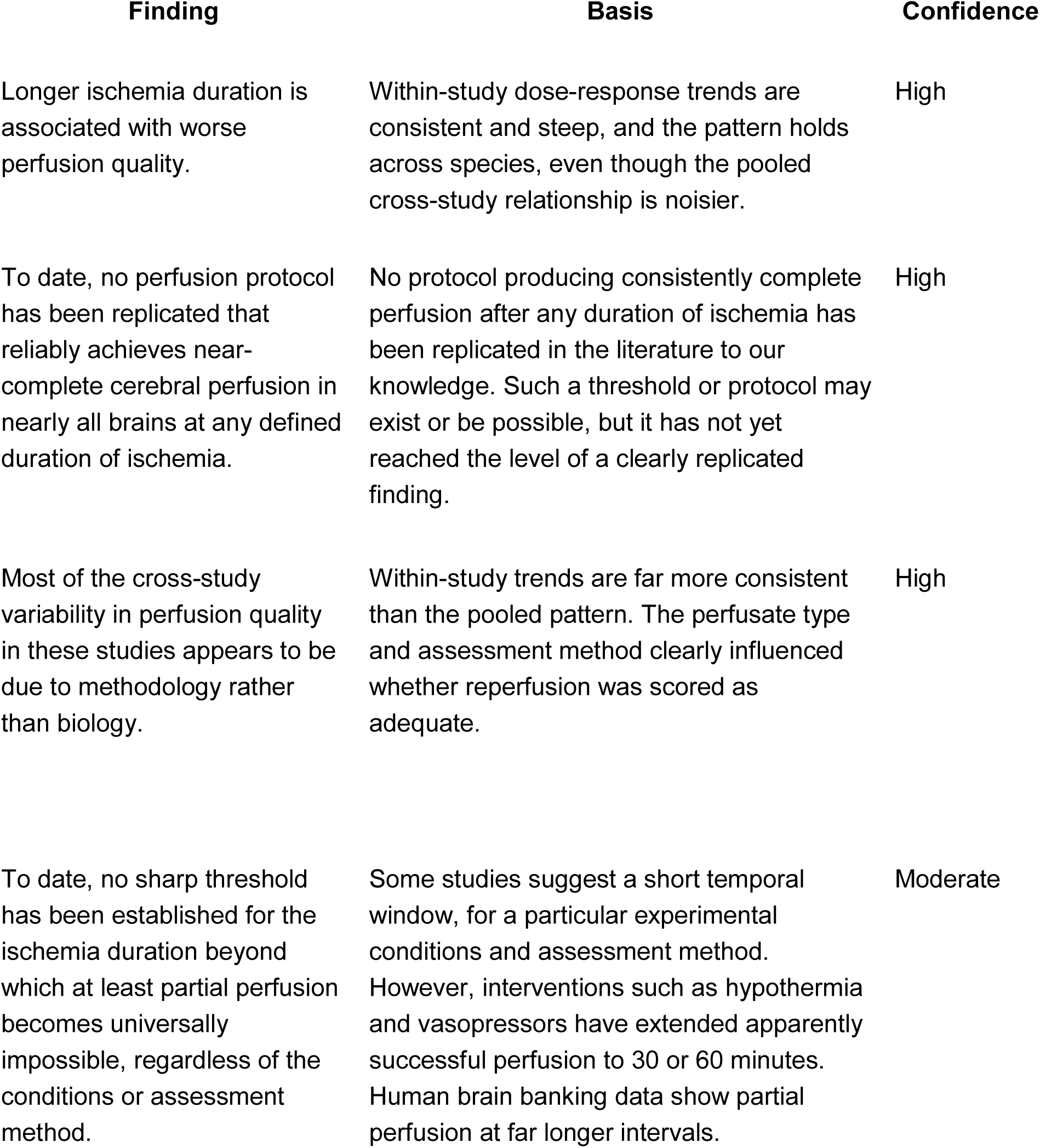
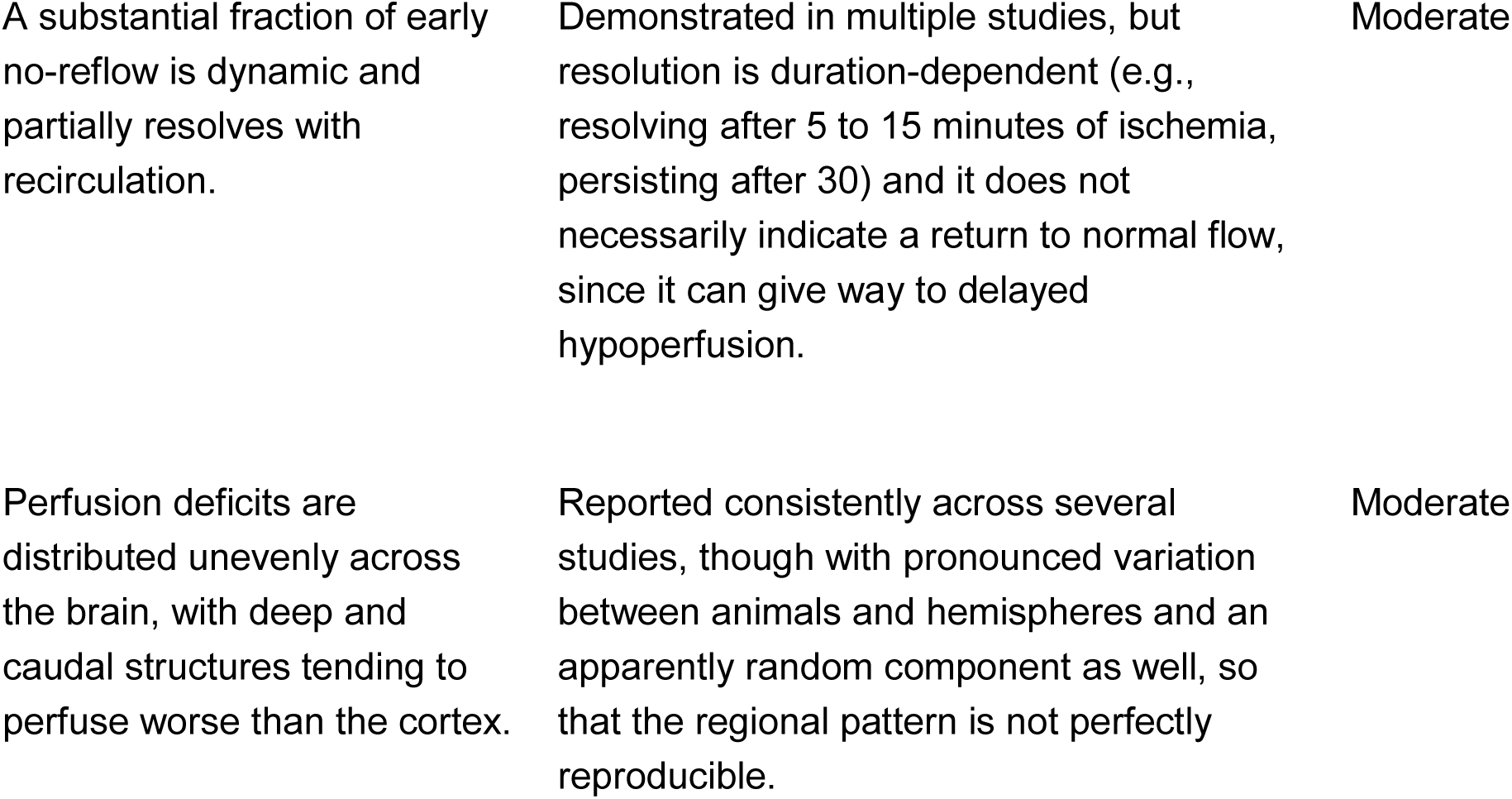
Several key findings of the review and the assessed confidence in each.

### Recommendations for future research

Future studies would benefit from a greater emphasis on prospective, preregistered experimental designs in which ischemia duration is systematically varied across multiple time points while the species, ischemia model, perfusion protocol, and assessment method are held constant. Such studies would provide a clearer picture of whether perfusion quality declines progressively with ischemia duration or instead exhibits one or more critical ischemia thresholds beyond which perfusion deteriorates rapidly. Detailed case-by-case reporting of perfusion outcomes, as exemplified by recent studies in structural brain preservation [17], would further improve reproducibility and facilitate comparison across laboratories. Independent replication of promising protocols should also be prioritized, as relatively few of the interventions reported to extend the apparent perfusability window have been independently reproduced.

Notably, the existing animal literature was generally not designed to identify the maximum duration of ischemia at which a given preservation protocol can still achieve tissue preservation indistinguishable from that obtained without ischemia. Indeed, our review suggests that there is unlikely to be a single universal perfusable window, because the answer depends on the experimental conditions, the perfusion protocol, and the endpoint used to define success. For example, a protocol that appears successful according to macroscopic or regional perfusion assessment methods may nevertheless have incomplete microvascular perfusion or otherwise fail to preserve ultrastructure at a level sufficient for neurite traceability by contemporary connectomics methods [3].

In human brain banking, some degree of postmortem ischemia is practically unavoidable. Carefully designed donation programs, with informed consent provided by the donors, may provide an opportunity to define the relationship between ischemia duration and preservation quality under realistic conditions in humans. Although this window may ultimately be extendable through pre-mortem or post-mortem interventions, implementing such approaches in human brain banking will require consideration not only of efficacy but also of practical, ethical, and legal constraints [35].

## Conclusions

The quality of brain specimens is crucial for understanding the mechanisms of neurologic and psychiatric disorders and improving our treatments for them. Our synthesis suggests that perfusion quality declines with ischemia duration, but that the timing and steepness of that decline depend heavily on the experimental context and on how perfusion quality is measured. We did not find evidence for an obvious threshold before which perfusion is nearly always complete and beyond which the brain becomes categorically unperfusable, regardless of such experimental and methodological context. This held for both measures we examined, i.e. the spatial extent of perfusion within the brain and the proportion of brains in a group perfused adequately. Several directions for future research follow. More studies that systematically manipulate the ischemia duration over more time points while holding the ischemia model, perfusion system, and assessment method constant would likely be the most informative way to characterize the decline. Because much of the cross-study variability appears to be methodological, the field would also benefit from more standardized and more sensitive measures of perfusion quality, ideally ones that can resolve the microvasculature across the brain rather than only confirming flow in a small number of brain regions. Furthermore, the interventions that extended the perfusable window in individual studies have rarely been replicated or compared head to head, so systematic testing of hypothermia, osmotic agents, and other interventions across different ischemic durations and species would help establish whether a reproducible protocol for consistent whole-brain perfusion after longer ischemia is achievable. Finally, given the relevance of this question to brain banking, forensic pathology, and other translational fields, more human data, particularly at the extended postmortem intervals typical in brain banking, would help bridge the gap between the animal literature and applied settings.

## Supplementary Data

**Data S1**. PRISMA 2020 checklist documenting adherence to the PRISMA reporting guidelines for systematic reviews.

**Data S2**. Title and abstract screening decisions exported from SysRev.

**Data S3**. Full-text screening decisions exported from SysRev.

**Data S4**. Final large language model (LLM) prompt used to extract data from the included studies.

**Data S5**. Validation of the LLM-assisted data extraction and study quality assessment. The workbook contains the original extracted data prior to any correction. Green cells indicate agreement with the manual review, yellow cells indicate non-substantive disagreements, and red cells indicate substantive disagreements.

**Data S6**. Final validated dataset used for analysis. The workbook contains the included studies, study quality assessments, and extracted data after manual review and correction of the validation subset where necessary. Studies included in the manual validation subset are highlighted in green.

**Data S7**. Interactive HTML version of Figure 2 (spatial completeness of brain perfusion after global ischemia), allowing zooming, panning, and display of study-level metadata by hovering over individual data points.

**Data S8**. Interactive HTML version of Figure 3 (frequency of adequate brain perfusion after global ischemia), allowing zooming, panning, and display of study-level metadata by hovering over individual data points.

## Abbreviations

CBF: Cerebral blood flow
CPR: Cardiopulmonary resuscitation
CT: Computed tomography
FITC: Fluorescein isothiocyanate
LLM: Large language model
PRISMA: Preferred Reporting Items for Systematic Reviews and Meta-Analyses.

## Author contributions

J.K., D.W., and A.T.M. conceptualized the study. J.K. and D.W. performed abstract screening, full text review, and data extraction. A.T.M. wrote the initial draft of the manuscript. B.W. critically reviewed the paper and contributed substantially to its content. All authors reviewed the manuscript and approved the final version.

## Acknowledgements

We would like to thank Brian Wowk for helpful personal communications regarding this topic.

## Declaration of Generative AI Technologies

In the preparation of this manuscript, the authors used Claude (Anthropic) and ChatGPT (OpenAI) for data extraction, coding assistance, and to improve the manuscript’s language. All AI tool-assisted content was reviewed and edited by the authors, who take full responsibility for the final publication.

## Data availability

R code and data used for analysis is available at https://github.com/andymckenzie/Ischemic_perfusion_review.

