## Supplementary material for "How long is the brain perfusable after global ischemia? A systematic review": DataS4_Data_Extraction_Prompt.pdf

### Data Extraction Prompt: Post-Ischemia Brain Perfusion Quality

You are extracting data from a research paper for a systematic review on how brain perfusion quality changes as a function of ischemia duration after global circulatory arrest (the "no-reflow phenomenon").

Read the paper fully, then output two tab-separated data rows inside code blocks --- one for Study Quality, one for Data Extraction (or multiple rows if there are multiple observations). These rows will be pasted directly into a spreadsheet that already has headers, so do not include column headers or labels inside the code blocks --- output only the data values, tab-separated, one row per line. The column order is specified below; match it exactly.

If a cell contains no data, leave it empty (two tabs in a row). If a cell would need to contain a tab or newline, use a space instead. **The contents of a cell must never include a tab character.** Tabs are the sole delimiter separating one cell from the next; this is what allows the output to be pasted directly into a spreadsheet. A tab appearing inside a cell's content will be misinterpreted as a column boundary, shifting that cell's remaining content and every subsequent cell one column to the right and corrupting the entire row. (See also the detailed "Tab prohibition within cells" rule in §1.3.)

Outside the code blocks, you may add any commentary, reasoning, or uncertainty notes in plain prose.

### §0 Purpose and Context

This section describes the goals of the review to help inform judgment calls during extraction. It does not override any specific instruction in later sections, but when a later section is ambiguous or does not address a situation, use this context to guide your decision.

**What the review is about.** We are building a dataset of every available empirical measurement of how well the brain can be perfused after varying durations of global cerebral ischemia. The central analysis is a plot of **ischemia duration** (x-axis) against **perfusion success** (y-axis), with observations annotated by species, perfusate type (blood vs. non-blood), treatment group (untreated vs. drug-treated), and other moderators. **Every distinct treatment group that underwent ischemia and had perfusion assessed contributes its own data point to this plot** --- treated and untreated groups are equally important as observations. Getting ischemia duration and the perfusion outcome right matters more than any other columns.

**The expected relationship** is a graded, stochastic decline in perfusion quality with increasing ischemia duration --- not a sharp threshold. There is substantial biological variability between individual animals at the same ischemia duration. This means borderline cases, partial perfusion, and mixed findings are valuable data points, not ambiguities to be cleaned up. Do not round uncertain findings into clean categories; preserve the nuance and let the analysis handle it.

**Who will use this.** The review serves researchers and practitioners in brain banking, biostasis/brain preservation, resuscitation science, organ transplantation, and forensic pathology. The unifying practical question across these fields is: **how long after circulatory arrest can you still deliver solutions through the brain's microvasculature?** Spatial completeness of perfusion (did perfusate reach all regions?) matters more than flow magnitude (how much perfusate reached each region?). Hyperperfusion is not a problem --- it means perfusate got through. Hypoperfusion and no-reflow (regions where perfusate cannot reach) are the outcomes of concern.

**What this means for extraction decisions:**

- When choosing which time point to extract from serial data, prioritize the **earliest** post-reperfusion measurement, because we want to characterize the initial post-ischemic state before secondary processes (resolution of no-reflow, delayed hypoperfusion) alter the picture.

- When a paper reports hyperemic CBF values, extract them faithfully. Hyperemia demonstrates that perfusion is achievable at that ischemia duration, which is informative.

- When a paper's findings are ambiguous between "adequate perfusion with some impairment" and "failed perfusion," err on the side of preserving the ambiguity in the data columns and explaining in the companion columns, rather than collapsing the finding into one category.

- Perfusate type (blood vs. non-blood) and species are key moderating variables. Be precise about these.

### §1 General Principles

These principles apply across the entire extraction process. Individual column definitions reference them by section number rather than restating them. When a column-specific instruction and a general principle appear to conflict, the column-specific instruction takes precedence for that column.

#### §1.1 Zero-Flow Boundary Identification

Many columns require identifying the precise boundaries of the ischemic (zero cerebral blood flow) period. The general rule: **count only the interval during which no blood reaches the brain.** Any period with partial circulation --- even inadequate circulation --- is not part of the zero-flow period.

This principle applies to: Ischemia Duration (column 9), Time from Ischemia Onset to Perfusion Quality Assessment (column 33), and Time from Perfusion Onset to Measurement (column 39). See §2 (Domain Knowledge) for specific applications to CPR, asphyxia, and DHCA models.

When computing any derived time value, state all component intervals explicitly and show the arithmetic in the companion Quote column. A single numeric value goes in the data column; all explanation goes in the companion.

#### §1.2 Companion Column Convention

The Data Extraction table has three structural sections:

**Identifier columns (1--4):** No companion columns.

**Paired extraction columns (5--34):** These alternate between a data column and its companion. In companion columns labeled "Quote," reproduce verbatim the passage from the paper that supports the data point, enclosed in quotation marks. If no supporting text exists, write "No supporting text." The companion column for Regional Breakdown is labeled "Source" --- record whether the regional data came from text, a specific table, or a specific figure.

**Critical rule:** Every paired data column must have its companion column populated, even when the data column is "N/A", "NR", or empty. For example, when Regional Breakdown is "N/A", Regional Breakdown Source must still be written as "N/A". Skipping a companion column drops a tab and shifts every subsequent column in the row.

**Derived/interpretive columns (35--44):** These alternate between a value column and a reasoning column.

**Terminal Notes columns (45--47):** Free-text, no companions.

#### §1.3 Data Column Formatting

Data columns (the non-Quote, non-Reasoning columns) must contain only the data value itself --- a number, a short categorical code, or a defined placeholder like "NR" or "N/A". All explanatory text, caveats, figure-estimation notes, and contextual details belong in the companion Quote or Reasoning column, or in Notes.

**Length constraint:** Data columns should almost never exceed 100 characters. If a data-column value is running longer than this, it probably contains explanatory text that belongs in the companion column instead. The exceptions are Regional Breakdown (column 29, which may list multiple regions) and Perfusate Type (column 17, which may describe multi-step protocols).

**Tab prohibition within cells:** Because the output format is tab-separated, a tab character inside any cell will be interpreted as a column boundary, pushing all subsequent content one column to the right and corrupting the entire row. **Never insert a tab character within the contents of any cell.** This applies to all columns in both the Study Quality and Data Extraction tables --- data columns, companion/Quote columns, Reasoning columns, and Notes columns alike. When you want to include multiple quoted passages, separate them with a space, semicolon, or the text string " | " --- never with a tab. When you want to indent or visually separate content within a cell, use spaces --- never tabs. Before finalizing each row, confirm that the only tab characters in the row are the delimiters between columns.

Examples of correct vs. incorrect formatting:

- Perfusion Pressure: write NR in the data column; put "No arterial pressure values reported; text states pressure was unchanged from baseline." in the Quote column. Do NOT write NR (systemic arterial pressure monitored but specific values not reported).
- Time from Ischemia Onset to Perfusion Quality Assessment: write 42 in the data column; put the arithmetic (e.g., "12 min ischemia + 30 min post-ischemia = 42 min") in the Quote column. Do NOT write 11-19 (8 min ischemia + ...).
- Sample Size: write a single number. If uncertain, write your best estimate; explain uncertainty in the Quote column. If the paper gives conflicting n values for the same group, use the lower number and note the discrepancy in the Quote column.

### §1.4 Data Source Hierarchy

When the same data point is available both in the running text (or a table) and in a figure, always extract from the text or table. Tag accordingly: "(from text)" or "(from Table N)". Use figure estimation only when the data point is not reported anywhere in the text or tables.

Never write "NR" if the value can be read or estimated from a figure. Before marking any column "NR," confirm that the value is genuinely not reported in the text, tables, *or* figures.

Figure-estimated values are less reliable than text-reported values and should always be flagged for manual review, but they are far more useful than "NR."

### §1.5 Figure-Estimation Protocol

When estimating a value from a figure, apply **two independent estimation methods** and compare them. This dual-estimation approach reduces the risk of large errors from any single visual judgment.

**Step 1 --- Identify the figure and state the basics:**

- State the figure number, the axis units (e.g., "x-axis is minutes post-ROSC; y-axis is ml/100g/min"), and the specific bar, point, or curve segment you are reading.

**Step 2 --- Method A (axis-referenced interpolation):** Identify the two nearest reference marks on the relevant axis (gridlines, tick marks, or labeled values) that bracket the target value --- one above and one below. State both reference values, then estimate where the target falls between them as a fraction (e.g., "approximately 1/3 of the way from the 20 to the 40 gridline"). Compute the interpolated value (e.g., 20 + 1/3 × 20 = 26.7). If the target falls below the lowest or above the highest reference mark, extrapolate and note this.

**Step 3 --- Method B (ratio to known reference):** Find a bar, point, or value in the **same figure** whose numeric value is known from the text or a table (e.g., a control group mean, a baseline value, or another group's reported result). Visually estimate the ratio of the target's height/position to this known reference (e.g., "the target bar appears approximately 60% as tall as the control bar"). Multiply the known value by this ratio to produce the estimate. If no text-reported reference value exists in the figure, use the most clearly readable bar or point as the reference value (estimated via Method A) and note that both values are figure-derived.

**Step 4 --- Compare and reconcile:**

- Compute the percent difference between the two estimates: |A − B| / ((A + B) / 2) × 100.
- If the two estimates agree within 20%: use their average as the final value. Append "(estimated from Fig. N, low confidence)".
- If the two estimates diverge by more than 20%: examine the figure again to identify the source of disagreement (e.g., ambiguous bar endpoint, overlapping error bars, compressed scale). Attempt to resolve the discrepancy. If it cannot be resolved, use the average but append "(estimated from Fig. N, very low confidence --- dual estimates diverged by [X]%)". The "very low confidence" flag signals that this value requires particularly careful manual verification.

**Step 5 --- Unit conversion:** Perform any required unit conversion explicitly (e.g., "ml/g/min × 100 = ml/100g/min").

**Recording:** In the companion Quote column, document both methods' reasoning in full: the reference marks used for Method A, the known reference value and visual ratio for Method B, both numeric estimates, the percent difference, and the final averaged value. In the data column, record only the final value with the ~ prefix.

**Flagging:** Prefix **all** figure-estimated values in data columns with "~" (e.g., "~27 ml/100g/min") so they are machine-identifiable for mandatory manual verification. Text- and table-derived values do not receive this prefix. All figure-estimated values are considered low confidence at minimum; the "very low confidence" flag is reserved for cases where the dual methods diverged by >20%.

#### §1.6 Unit Conversion

When computing any derived value, verify that all component values are in the same units before performing arithmetic. If conversion is needed, show the conversion explicitly. Common pitfalls: adding minutes to hours without converting (12 min + 0.5 hours ≠ 12.5; it equals 42 min), confusing ml/g/min with ml/100g/min (factor of 100), mixing mmHg with $cmH_2O$ or torr.

#### §1.7 Self-Check Requirements

**Tab counting:** After generating each tab-separated row, count the number of tab characters. The Study Quality table has 17 columns (16 tabs per row). The Data Extraction table has 47 columns (46 tabs per row). If your count differs, identify and fix the misalignment before outputting.

**Spot-check columns:** After generating each Data Extraction row, verify these columns by their 1-indexed column number:

- Column 4 (Additional Outcome Specifier): must be blank or a short label (e.g., "untreated", "rt-PA treated", "20 min ischemia"). If it contains a species name or quote text, you have a tab error in columns 1--4.

- Column 30 (Regional Breakdown Source): must be one of: "from text", "from Table N", "estimated from Fig. N (low confidence)", "estimated from Fig. N (very low confidence)", or "N/A". If it contains a brain region name, you have dropped a tab.

- Column 35 (Blood vs. Non-Blood): must be exactly "Blood", "Non-Blood", or "Mixed".

- Column 41 (Perfusion Quality Category): must be exactly one of the seven defined categories.

**Verification line:** After outputting each Data Extraction row, immediately output a verification line in the following format (outside the code block):

```
VERIFY ROW: col35=[value] | col41=[value]
```

where [value] is the exact content you placed in that column. This provides a machine-checkable confirmation that columns 35 and 41 contain valid categorical values and have not been displaced by a tab error.

**Content-type check:** After generating each row, scan every data column (non-Quote, non-Reasoning). If any data column contains more than ~100 characters of text, a complete sentence, or quotation marks, it almost certainly contains companion-column content that has leaked into a data column due to a tab error. Fix before outputting.

**Completeness audit (after all rows are generated):** Compare your final set of rows against the group inventory from §3.0. Confirm that every non-sham group in every data table has a corresponding row. If any treatment group's perfusion data appears only in a Notes column rather than in its own row, this is an extraction error --- create the missing row before finalizing.

If any spot-check fails, there is a column shift. Fix it before outputting. **Do not trust a row that has not passed the anchor-cell check in §1.7A.** A row with correct biological content but wrong column placement is considered incorrect.

### §1.7A Anchor-Cell Output Lock

Before outputting each final Data Extraction row, perform this mechanical check in order:

1. Build the row as a numbered list of **47 cells** first, not as raw TSV.
2. Confirm these anchor cells before converting to TSV:

- **30 = Regional Breakdown Source**
- **31 = Brain Region(s) Assessed**
- **33 = Time from Ischemia Onset to Perfusion Quality Assessment (numeric only)**
- **35 = Blood / Non-Blood**
- **37 = Caveats Category**
- **39 = Time from Perfusion Onset to Measurement (numeric only)**
- **41 = Perfusion Quality Category**
- **43 = Converted % Brain Perfused**

3. Then convert the numbered list to a single TSV row.

4. After conversion, re-check that:

- column 30 is a source code, not a sentence,
- columns 33 and 39 are numeric or "NR",
- column 35 is exactly "Blood", "Non-Blood", or "Mixed",
- column 41 is exactly one of the allowed category labels,
- column 43 is a number, "Cannot convert", or "NR" only.

5. If any of these checks fail, rebuild the row before outputting.

**Special warning:** The most common row-corruption failure is dropping column 30 when Regional Breakdown is "N/A". If column 29 is "N/A", column 30 must still be exactly "N/A".

### §1.8 No-Reflow Priority Rule

No-reflow is the core phenomenon of this review. When categorizing perfusion quality (column 41):

1. If no-reflow is present in any region → "No-reflow present (quantified)", regardless of what other regions show.
2. If no-reflow was assessed and found absent → "No-reflow absent", regardless of whether the dominant finding is hyperemia or hypoperfusion.
3. Use "Hypoperfusion" or "Hyperperfusion" ONLY when the study did not assess no-reflow at all (e.g., measured only CBF without any spatial or tracer-based assessment of vascular patency).
4. Use "Mixed regional heterogeneity" only when no-reflow was not assessed and the CBF pattern does not fit a single category.

Before writing "Hypoperfusion" or "Hyperperfusion," confirm that the paper does not report any spatial perfusion assessment that could indicate whether no-reflow was present or absent. **Multi-region CBF methods** (microspheres counted across multiple brain regions, multi-region hydrogen clearance, autoradiography) provide spatial resolution: if a region received zero flow, the method would detect it. When such a method finds measurable flow above zero in all assessed regions, this constitutes sufficient spatial evidence for "No-reflow absent" --- even if the dominant pattern is hyperemia. Record the flow magnitude (including hyperemia) in the

Perfusion Outcome column; the Perfusion Quality Category captures only the spatial patency finding.

**Qualitative whole-brain methods** (fluoroscopic angiography, gross tracer filling inspection) can visualize the brain's entire vascular territory but lack the resolution to detect capillary-level no-reflow. No-reflow in this literature is predominantly a microvascular phenomenon (capillary plugging, endothelial swelling, microthrombi) at a spatial scale below what these methods resolve. A brain can show "robust filling" of all major arteries on angiography while having substantial capillary-level no-reflow throughout. Therefore:

- These methods **can** support "Qualitative flow present" or "Qualitative flow absent."
- These methods **can** support "No-reflow present (quantified)" if they show absent filling in a vascular territory (large-vessel obstruction guarantees absence of downstream microvascular flow).
- These methods **cannot** support "No-reflow absent," which requires a method with microvascular or tissue-level resolution (microspheres, autoradiography, tracer filling studies, histological capillary patency assessment).

**Hyperemic data:** Consistent with the review's goals (§0), hyperperfusion is not a concern --- it indicates perfusate can reach tissue. Extract and report hyperemic data faithfully, including readings from the transient hyperemic phase. These demonstrate that perfusion is achievable at a given ischemia duration.

This rule applies regardless of the suspected mechanism of perfusion failure: even if authors attribute near-zero flow to dynamic vasospasm rather than fixed capillary obstruction, categorize as "No-reflow present (quantified)" if the spatial assessment shows regions with absent or near-absent flow. The mechanistic distinction is captured in the Reasoning column (column 42), not the category.

**Mechanism-independence extends to §3.2 transition rows.** The same mechanism-independence principle that governs categorization also governs transition-row decisions. Once a measurement has been assigned a Perfusion Quality Category under §1.8 --- regardless of the suspected mechanism --- any subsequent shift to a different category at a later time point is evaluated under §3.2 based solely on the category labels, not on the extractor's assessment of what caused the category to change. Confounding variables that may explain the transition (temperature normalization during DHCA rewarming, perfusion pressure changes due to vasopressor support, $PaCO_2$ shifts, hematocrit changes, anesthetic depth changes, etc.) are documented in the Perfusion Quality Category Reasoning column (42), column 46 (Notes - Long-Term Perfusion Improvement), and/or columns 37--38 (Caveats), but do not affect row-creation decisions. Applying a mechanism-based exception to suppress a transition row while retaining the mechanism-independent categorization that created it is an internal inconsistency --- the categorization and its downstream consequences must use the same logic.

#### §1.9 Spatial Completeness vs. Flow Magnitude

The Converted % Brain Perfused column (43) measures **spatial completeness** --- what fraction of brain tissue received any blood flow at all --- not flow magnitude relative to baseline.

- Flow at 60% of baseline but present everywhere → 100% (the entire brain is spatially perfused).
- 30% of brain has zero flow → 70%, regardless of flow magnitude in perfused regions.

This distinction also informs the % Brains Adequately Perfused column (27): when a spatially resolved method (microspheres, autoradiography, multi-region hydrogen clearance) finds measurable flow above zero in all assessed regions, infer 100% adequately perfused using "measurable CBF in all regions" as the threshold.

**Spatial resolution requirement.** Both columns 27 and 43 ask spatial questions --- they require knowing whether flow reached different parts of the brain, not just whether flow was present at one location or a subset of locations. Methods whose field of view does not encompass the whole brain cannot answer these questions, regardless of sample size or whether all animals show measurable flow. This includes single-site methods (one laser Doppler probe, one thermocouple, single-site laser speckle imaging through a cranial window) and methods with spatial resolution over only part of the brain (SDF/OPS imaging of a cortical patch, cranial window preparations covering one region). For such methods, write "NR" in both columns 27 and 43, and note in the companion column that the method's field of view does not cover the whole brain. The actual flow values from these methods are still fully captured in column 23 (Perfusion Outcome), which is where they belong.

**Whole-brain coverage clarification.** Some methods sample many regions but still do **not** cover the whole brain. Multi-slice imaging of only a few coronal levels, partial-field CT/MRI coverage, or a restricted set of ROIs does **not** justify whole-brain conversion unless the paper explicitly states that the sampled regions constitute the whole brain or provides a true whole-brain/global spatial assessment. Example: three Xe/CT coronal slices with many ROIs = **not** whole-brain coverage for columns 27 and 43.

**Group-level whole-brain inference clarification.** If a spatially resolved method samples named regions spanning the major anatomical compartments of the brain **and** either (a) the paper explicitly states that there were no regional differences or that all assessed regions had measurable flow, or (b) tabulated or graphed data show all assessed regions with nonzero mean flow values, you may infer 100% for columns 27 and 43 **only when the method genuinely covers the whole brain rather than a subset of slices or a local field of view.** State the inference basis explicitly in the companion column. Condition (b) ensures that microsphere studies and similar multi-region methods where whole-brain patency is evident from the data are not marked "NR" solely because the authors did not write an explicit summary statement.

#### §1.10 Data Source Audit

Before completing each row, check all tables and figures in the paper for data that might populate empty columns. If a column is marked "NR," confirm that the value is genuinely not reported anywhere --- including hemodynamic tables, figure axes and legends (which may state variability type, sample sizes, or units), supplementary materials, and **protocol descriptions** (which may state target perfusion pressures, e.g., "MAP was maintained at >80 mmHg"). A target pressure stated in the methods section is a reportable perfusion pressure value --- record it with the appropriate qualifier (e.g., ">80 (MABP, target)") and note in the Quote column that it is a protocol target rather than a measured group mean. **However, if the protocol target was maintained only transiently (e.g., for the initial 15 min of recirculation) and the perfusion quality measurement occurs later (e.g., at 30 min), note this temporal mismatch in the Quote column and append "(transient, may not reflect pressure at measurement time)" to the data value.** When the pressure at the actual measurement time is unknown, this qualifier alerts downstream analysis that the value may not represent conditions during assessment.

#### §1.11 Variability Type Identification

When extracting variability data (column 25), always quote the paper's exact descriptor for the variability measure (e.g., "± S.E.M.", "± SD", "95% CI", "± SE") in the companion Quote column. Do not infer the type from context or convention --- if the paper uses "±" without specifying the type anywhere in the text, methods, or figure legends, write "SD/SEM unspecified." Check the methods section, figure legends, and table footnotes for declarations like "values are means ± SEM" that may apply globally to all reported data.

#### §1.12 Caveats Classification

Column 37 flags experimental conditions that differ from the baseline scenario relevant to this review. Apply §1.12 when populating column 37.

**Important:** Caveats are annotations applied to each row's own data. Flagging a treatment as a caveat does **not** replace the requirement to create a separate row for that treatment group (§3.5). Each treatment group gets its own row with its own Caveats Category.

**Guiding principle.** This review serves human brain banking, biostasis, and related fields where the practical question is: how well can the brain be perfused after circulatory arrest under conditions typical of human death? The caveats system flags experimental conditions that would not be present in a human postmortem context, so that downstream analysis can assess how much the experimental literature's perfusion results depend on interventions unavailable in practice.

**Categories** (select all that apply, semicolon-separated):

- **None beyond standard resuscitation:** Use only when reperfusion occurs spontaneously or via a simple procedure (clamp release, defibrillation, single standard-dose epinephrine bolus during CPR) without ongoing pharmacological support or other interventions during the measurement period. See "Standard resuscitation definition" below.

- **Pre-treatment (experimental intervention):** Any drug, agent, or manipulation administered before ischemia that could plausibly affect post-ischemic perfusion quality and would not be present in a typical human postmortem scenario (e.g., a neuroprotective agent, vasodilator, or thrombolytic given pre-arrest).

- **Pre-treatment (standard experimental anticoagulation):** Heparin or other anticoagulants administered before ischemia as part of standard experimental protocol. Flagged separately because anticoagulation reduces clot-mediated no-reflow and is not typically present in human death, but is near-universal in experimental studies --- particularly those using microspheres.

- **Intervention to improve perfusion quality:** Any drug, agent, or non-pharmacological intervention applied during reperfusion or assessment intended to improve perfusion (e.g., thrombolytics, vasodilators, therapeutic hypothermia, hemodilution).

- **Intervention that worsens perfusion:** Any agent applied during reperfusion expected to worsen perfusion (e.g., NO synthase inhibitors as mechanistic probes).

- **Post-resuscitation vasopressors:** Any vasopressor or inotrope administered during the reperfusion period to maintain or elevate blood pressure, including continuous infusions of epinephrine or norepinephrine to maintain target MAP. This is distinct from a single standard-dose epinephrine bolus during CPR (see below).

- **Hypothermic ischemia:** Ischemia conducted under hypothermic conditions.

- **Other (specify):** When in doubt about classification, use this and describe.

**Decision rules:**

1. *Dual-phase interventions:* If a drug is initiated before ischemia AND continued during reperfusion, flag both "Pre-treatment" (experimental intervention or standard experimental anticoagulation, as appropriate) and "Intervention to improve perfusion quality." Example: a 10 mg/kg bolus of Drug X before arrest followed by 5 mg/kg/hr infusion during reperfusion gets both flags.

2. *Standard resuscitation definition:* A single epinephrine bolus at ≤20 μg/kg during CPR is standard (roughly equivalent to AHA-recommended human dosing scaled by body weight). Defibrillation and mechanical ventilation are standard. Everything beyond this

requires a flag:

- continuous epinephrine or norepinephrine infusions → "Post-resuscitation vasopressors"
- repeated epinephrine boluses beyond the initial one → "Post-resuscitation vasopressors"
- a **single high-dose epinephrine bolus >20 μg/kg during CPR** → "Other (specify: high-dose CPR epinephrine)"
- any drug given to modify perfusion → "Intervention to improve perfusion quality"

If a study includes both a high-dose CPR epinephrine bolus and a later vasopressor infusion, apply both relevant flags.

3. *Post-hoc subgroups:* Groups split by functional recovery or other achieved variables are handled by labeling in column 4, not by a Caveats flag (see §3.8).
4. *Routine model-management procedures:* Hemodynamic management steps that are inherent to the experimental model and aimed at maintaining physiological parameters within normal ranges are not caveats, unless they plausibly augment cerebral perfusion above what the standard resuscitation protocol would achieve. Examples of non-flaggable procedures: temporary partial occlusion of a distal vessel during the surgical transition from clamp release to spontaneous circulation (to prevent systemic hypotension), conditional blood withdrawal during ischemia to prevent hypertensive crisis, and reinfusion of withdrawn blood after ischemia. These are safety or model-management necessities, not perfusion-enhancing interventions. If the same procedure also has a plausible perfusion-augmenting effect (e.g., temporarily redirecting a disproportionate fraction of cardiac output to the brain), note this in the Caveats Details column (38) or Notes (45) without assigning a Caveats Category flag. However, if a model-management procedure incidentally introduces a perfusion-modifying agent into the circulation (e.g., withdrawn blood is heparinized to prevent clotting in the syringe, then reinfused), the *procedure* is exempt but the *agent* still requires its appropriate Caveats Category flag (in this example, "Pre-treatment (standard experimental anticoagulation)").

### §2 Domain Knowledge

These are factual points that inform extraction decisions. They are stable across papers.

#### §2.1 CPR and Partial Circulation

Cardiopulmonary resuscitation (CPR), including chest compressions and cardiac massage, produces partial cerebral circulation. The CPR period is therefore NOT part of the zero-flow ischemic interval (§1.1). For cardiac arrest models with CPR:

- **Ischemia duration** (column 9) = the interval before CPR began (e.g., the VF-only period, or the interval from arrest induction to start of chest compressions). CPR produces partial cerebral circulation, so it is not ischemic time.

- **Time from ischemia onset to perfusion quality assessment** (column 33) = total elapsed time from ischemia onset to the moment perfusion quality is measured. This is not the time to the start of reperfusion --- it is the time to the assessment. For cardiac arrest with CPR, this equals: ischemia duration + CPR duration + post-ROSC interval to measurement. CPR time is neither ischemia (column 9) nor confirmed full perfusion (column 39); it occupies an intermediate zone where partial circulation of uncertain magnitude occurs. It is captured implicitly as part of column 33 but not counted in either column 9 or column 39.

- **Time from perfusion onset to measurement** (column 39) = post-ROSC interval only (time from ROSC to assessment). ROSC marks the start of confirmed perfusion.

In the Quote column for column 33, list all component intervals explicitly using this format: "VF: X min + CPR: Y min + post-ROSC: Z min = [total] min."

*Example 1:* A study induces 8 min of VF, followed by 8 min of CPR before ROSC, then measures perfusion at 2 h post-ROSC. Column 9 = 8 min. Column 33 = 8 + 8 + 120 = 136 min. Column 39 = 120 min.

*Example 2:* A study induces 15 min of VF, followed by 12 min of CPR before ROSC, then measures perfusion at 30 min post-ROSC. Column 9 = 15 min. Column 33 = 15 + 12 + 30 = 57 min. Column 39 = 30 min.

When a paper provides separate ranges for total arrest duration (VF + CPR) and VF-only duration, always use the VF-only range as the basis for ischemia duration.

#### §2.2 Asphyxia Models

In asphyxia cardiac arrest models, cardiac arrest may begin partway through the asphyxia period. Report the protocol duration as ischemia duration and note the discrepancy if the actual zero-flow period is shorter.

#### §2.3 Deep Hypothermic Circulatory Arrest (DHCA)

For DHCA followed by rewarming, the start of antegrade cardiopulmonary bypass (CPB) rewarming marks the beginning of reperfusion. Extract the earliest rewarming measurement as the primary observation, even if perfusion impairment at that point may partly reflect incomplete rewarming physiology rather than ischemia-induced vascular obstruction. Note potential confounding by hypothermic physiology in the Perfusion Quality Category Reasoning column (42).

**DHCA rewarming and §3.2 transition rows:** If the Perfusion Quality Category (column 41) changes between the earliest rewarming measurement and a later measurement --- e.g., from "No-reflow present (quantified)" during early rewarming to "No-reflow absent" at end of rewarming --- this constitutes a qualitative state change under §3.2 and requires a transition row, even though the perfusion recovery coincides with temperature normalization. See §1.8 (mechanism-independence extends to §3.2 transition rows). Document the temperature confound and the degree to which the observed no-reflow may reflect hypothermic physiology versus ischemia-induced obstruction in column 42 and column 46. The "Hypothermic ischemia" caveat flag (column 37) enables downstream sensitivity analysis to handle these observations appropriately.

#### §2.4 Figure Time Axis Convention

When a figure axis is labeled "time following [occlusion / arrest / ischemia]," interpret this as time from the start of the ischemic insult (i.e., axis zero = ischemia onset). Do not add the ischemia duration on top --- that would double-count the ischemic period.

#### §2.5 Ischemia Model Eligibility

The ischemia must be both **complete** (zero blood flow to the brain) and **global** (affecting the entire brain). These two criteria are independent.

**Eligible models** (complete and global by design):

- Cardiac arrest (any induction: KCl, electrical fibrillation, asphyxia, exsanguination, etc.)
- Ventricular fibrillation (hemodynamically equivalent to cardiac arrest)
- Decapitation
- Raised intracranial pressure above mean arterial pressure (if ICP genuinely exceeds MAP)

**Models requiring careful judgment** (may or may not qualify --- read the paper closely):

- Aortic occlusion / intrathoracic vascular clamping: if the clamp is proximal to the brachiocephalic trunk, should block all forward flow, but retrograde collateral flow is theoretically possible. Ascending aorta occlusion is sometimes performed alone and sometimes combined with inferior vena cava (IVC) occlusion to prevent venous return from maintaining residual cardiac output. These are different protocols with different confidence levels for complete ischemia. Include only if the paper provides no evidence of residual cerebral blood flow.

- Neck tourniquet / neck cuff compression: completeness depends on pressure and anatomy. Include only if the paper demonstrates or clearly claims zero residual CBF.

For any model in this category, state in Quality Notes: "Completeness confidence: High/Medium/Low" with a brief justification. Give substantial weight to empirical verification: if the study measured CBF during ischemia and excluded animals with residual flow above a threshold (e.g., >5 ml/min/100g), this supports a "High" confidence rating even if the model could theoretically permit residual flow. For well-established models (cardiac arrest, VF, ascending aorta occlusion with IVC occlusion), "High" may be based on the broader literature, provided the paper does not report evidence of residual flow.

**Model description specificity (column 13):** For vascular occlusion models, record all vessels occluded (e.g., "ascending aorta occlusion with IVC occlusion," not just "aortic occlusion"). Different occlusion configurations may differ in whether they achieve complete ischemia, so the specific protocol must be identifiable from the extraction.

**Excluded models:**

- Four-vessel occlusion (4-VO): residual collateral flow through spinal, muscular, or anastomotic pathways is common. Studies frequently accept up to 25% residual CBF.

- Bilateral carotid occlusion alone: leaves vertebrobasilar circulation intact. Not global.

- Forebrain ischemia models (e.g., bilateral carotid occlusion in gerbils lacking a complete circle of Willis). Not global.

- Focal/regional models (e.g., middle cerebral artery occlusion). Not global.

- Any model where the paper reports or accepts residual CBF during ischemia. Not complete. **Exception:** if the paper reports very low measured CBF (e.g., <1--2 ml/100g/min) during ischemia but the authors explicitly attribute this to measurement artifact (tracer evaporation, extracerebral radioactivity, skin blood flow contamination) rather than actual perfusion, the paper is not excluded. Note the reported value and artifact attribution in Quality Notes.

**Retrograde cerebral perfusion (RCP) groups:** Extract only the standard circulatory arrest group. RCP represents attempted perfusion via an alternative route during the arrest period; capillary perfusion failure during RCP reflects hemodynamic inadequacy of retrograde delivery, not ischemia-induced no-reflow. RCP also introduces confounding factors (venous hypertension, brain edema) qualitatively different from post-ischemic insult. Note the RCP group's existence and key findings in the Notes column of the standard arrest group's row.

**Other exclusions:** Reviews/editorials without original data, case reports, articles not available in English.

If the paper is excluded, state the reason at the top of your response and still output the Study Quality row so the exclusion can be documented.

### §2.6 What Qualifies as Perfusion Quality Data

Any empirical measure of how well the brain can be perfused post-ischemia: tracer distribution (carbon black, India ink, Evans blue, FITC-dextran, fluorescein, HRP used as a perfusion tracer), CBF measurements (microspheres, $^{14}$C-iodoantipyrine autoradiography, $^{133}$Xe clearance, hydrogen clearance, laser Doppler flowmetry, ASL-MRI, Xe-CT, PET-based CBF, laser speckle imaging), TTC staining, histological grading of vascular patency, angiography, FITC-albumin fluorescence microscopy, etc. HRP used solely as a BBB permeability marker does not qualify. Post-ischemic CBF data collected incidentally (e.g., as a covariate) qualifies if usable. However, CBF measured as part of a pharmacological dose-response experiment (e.g., at multiple induced CPP or $PaCO_2$ levels) is not a standard perfusion quality observation and is handled under §3.9, not extracted as a primary row.

**Quantitative or semi-quantitative requirement.** The measure must produce a numeric value, a score on a defined scale, a percentage, a spatial map, or at minimum a binary per-animal classification with group-level counts. Representative images alone (e.g., a single fluoroscopic angiogram per group) do not qualify unless the paper also reports explicit per-animal tallies or a group-level statement about the consistency of the finding (e.g., "flow was present in 6/6 OrganEx vs. 0/6 ECMO"). If such a group-level statement exists and the sample size is known, infer per-animal counts and note the inference in the companion Quote column.

**Binned histogram data** qualifies: if the paper reports the number of animals in defined perfusion-quality bins (e.g., "13/46 <20 ml/100g/min, 22/46 20--40, 11/46 >40"), record the distribution in Perfusion Outcome and convert to % Brains Adequately Perfused using a threshold derived from the paper's own definition. If no threshold is defined, use the most stringent bin boundary and note this choice.

**Non-qualifying measures:**

- Purely narrative/qualitative descriptions of vascular patency without any numeric scoring or grading.

- Circular designs where animals are sorted by perfusion status and then perfusion status is reported as an outcome. This includes post-hoc grouping by recirculation quality --- e.g., a study that divides animals into "impaired recirculation" and "unimpaired recirculation" groups based on pial vessel observation and then reports perfusion-related morphological findings per group. Extracting such groups would guarantee the reported association by construction, contributing no information about the probability of perfusion failure at a given ischemia duration. If merging the post-hoc groups back into a single unsorted cohort is possible (i.e., individual-animal ischemia durations can be paired with individual perfusion outcomes), do so and extract accordingly. If the paper reports only group-level summaries with overlapping ischemia duration ranges, the data cannot be deconvolved and should not be extracted as primary rows. Note the group-level findings in column 45 (Notes) of any extractable row from the same paper, or in the Study Quality Notes if no rows are extractable.

- Tissue viability stains (e.g., TTC) performed many hours after reperfusion: these reflect cumulative ischemia--reperfusion cell death, not the initial quality of post-ischemic perfusion.

- Single-site tissue oxygenation monitors (e.g., INVOS/NIRS cerebral oximetry): these measure oxygen saturation at one cortical location, not cerebral blood flow, vascular patency, or spatial perfusion distribution.

- Single-path or single-site hemodynamic kinetics: methods that measure only the *speed* or *transit time* of flow through a single vascular path, a single artery-vein pair, or a single measurement site --- without characterizing spatial perfusion distribution across the brain --- do not independently qualify as perfusion quality data. Examples include: mean transit time (MTT) measured from a single pial artery to a single pial vein through one cortical field, single-probe thermodilution transit time, and single-site bolus-tracking kinetics. These methods characterize how *quickly* perfusate moves through tissue that is already reached, not *where* perfusate reached or failed to reach, and therefore do not map onto the review's core spatial-patency question (§0). Document such findings in the Notes column (45) of the row for the qualifying perfusion modality from the same paper. This exclusion does not apply to transit-time methods with whole-brain or multi-regional spatial coverage (e.g., whole-brain CT perfusion MTT maps, multi-slice bolus-tracking MRI), which do provide spatial information and qualify normally.

**Hypothermic studies:** Include but flag in Notes for sensitivity analysis.

**Companion paper references.** When a paper explicitly declines to present its perfusion-relevant data, instead deferring to a separate publication (e.g., "the physiological findings have been presented elsewhere [citation]," or "the hemodynamic data are reported in [citation]"), note the cited companion paper's full reference (authors, year, journal, volume, pages) in Quality Notes and flag it for inclusion screening if it is not already in the review's corpus. Complete the extraction of whatever perfusion quality data is independently available in the current paper, but do not attempt to extract data that the paper attributes to the companion publication. If the current paper contains no independently extractable perfusion quality data beyond what it attributes to the companion, this is not grounds for exclusion --- output the Study Quality row, note in Quality Notes that the perfusion data resides in the companion paper, and produce no Data Extraction rows. **Important:** This rule applies only when the current paper defers to the companion --- i.e., it explicitly states the data was presented or reported in the other publication and does not reproduce it. If the current paper presents the data as its own original findings, extract it normally, even if a companion paper also reports the same data. The goal is to avoid extracting data the paper itself says it is not presenting, not to deduplicate across papers.

#### §2.7 Time-Window Averages

When a paper reports perfusion outcome as an arithmetic average over a time window (e.g., "mean CBF over 5--30 min post-ROSC") rather than at a discrete time point, use the **midpoint** of the window as the time value for columns 33 and 39. In the companion Quote column, state the full window (e.g., "Reported as average over 5--30 min post-ROSC; midpoint = 17.5 min used"). Do not use the window start as the time value --- the reported measurement does not represent conditions at the start of the window alone. If a discrete earliest time point is also reported separately, prefer that value over the window average per §3.2.

**Exponential washout methods ($H_2$ clearance, inert gas clearance):** These methods derive a rate constant (K) from the exponential decay of a tracer signal over a multi-minute washout period. Although the washout takes many minutes, the derived K value is **not** an arithmetic time-window average --- it is an exponential fit dominated by the initial slope of the washout curve. Do not apply the midpoint rule above to these methods. Instead, treat the moment of assessment as the **start** of the washout recording (i.e., the moment the tracer-free gas mixture begins delivery or, equivalently, the moment perfusion-driven washout begins). In the companion Quote column, note the total washout duration and that the CBF value reflects an exponential rate constant weighted toward the early washout period.

### §3 Observation Rules

An observation is one row in the Data Extraction table. These rules govern how many rows a paper generates.

> ⚠ **MOST COMMON EXTRACTION ERROR:** Failing to create separate rows for treatment groups. Every group that underwent ischemia and had perfusion assessed gets its own row --- including drug-treated groups, even when n=1. Do

NOT summarize treatment group data in the Notes columns of the untreated row. See §3.5.

⚠ **ROW-COUNT TIE-BREAKING RULE:** When the rules in §3 leave genuine ambiguity about whether a situation warrants one row or two (or more), always err on the side of creating more rows, not fewer. An extra row that turns out to be unnecessary can be removed during manual review with no data loss. A missing row represents lost data that may never be recovered, because the reviewer would need to re-read the original paper to discover the omission. This asymmetry means the cost of under-extraction always exceeds the cost of over-extraction. If you find yourself constructing an argument for why a row is not needed, treat that reasoning itself as a signal that the row should probably be created --- document your uncertainty in the companion columns and let the downstream analysis decide.

### §3.0 Pre-Extraction Group Enumeration (Required)

Before writing any data rows, list every extractable experimental group in the paper as a numbered inventory. For each group, state: ischemia duration, treatment (or "untreated"), assessment timing, and expected sample size. **For case-series papers with serial measurements per animal (§3.2A), also state for each animal whether a qualitative state change (§3.2) is present; if so, name the two time points that define the transition and list the transition row in the inventory.** Then state the total number of rows you expect to produce. **You are committed to this row count — do not reduce it during row construction.**

*Example 1:* "Groups identified: (1) 5 min untreated, immediate, n=3; (2) 5 min heparin 20 mg/kg, immediate, n=2; (3) 10 min untreated, delayed 30 min, n=4; (4) 10 min heparin 20 mg/kg, delayed 30 min, n=1. Expected rows: 4."

*Example 2 (case-series with transitions):* "Groups identified: (1) Animal A, 10 min VF, earliest at 3 min post-ROSC, n=1 — qualitative state change present (no-reflow absent → present, transition at 28 min); (2) Animal A transition row, 28 min post-ROSC; (3) Animal B, 10 min VF, earliest at 27 min post-ROSC, n=1 — qualitative state change present (no-reflow absent → present, transition at 48 min); (4) Animal B transition row, 48 min post-ROSC; (5) Animal C, 10 min VF, earliest at 18 min post-ROSC, n=1 — qualitative state change present (no-reflow absent → present, transition at 50 min); (6) Animal C transition row, 50 min post-ROSC. Expected rows: 6."

**A small case-series study may produce more rows than its sample size — e.g., 6 rows from 3 animals. This is correct and expected.** Row count reflects the number of distinct observations, not the statistical weight of the study. If you find yourself writing "transition data documented in column 46 for brevity" or reducing the row count after committing to it, this is the extraction error the consistency requirement (§3.2) exists to prevent.

After completing all rows, verify that every group and transition in your inventory has a corresponding row. If any is missing, add it before finalizing.

### §3.1 Core Rule

One row per distinct combination of **experimental group** and **assessment time**. An experimental group is a set of animals that received the same ischemia duration, same ischemia model, and same treatment.

### §3.2 Multiple Time Points for the Same Experimental Condition

When a paper reports perfusion measurements at multiple post-reperfusion time points for the same experimental condition --- whether using the **same set of animals** measured serially (repeated measures) or **separate cohorts of animals** each assessed at a single time point (parallel-cohort design) --- extract only the **single most representative time point** per experimental condition. Choose the **earliest** post-reperfusion time point that captures the initial perfusion quality finding. This review is interested in how well the brain can be perfused shortly after ischemia, so the initial post-ischemic state takes priority over later states where impairment may have resolved.

**Exception --- qualitative state change:** If the serial data show a qualitative transition in **spatial perfusion patency** (e.g., no-reflow absent at 5 minutes evolving into no-reflow present by 30 minutes, or no-reflow at 5 minutes resolving to full patency by 30 minutes), extract **two** rows: one for the earliest time point and one for the time point at which the qualitative change is first evident. Use column 4 (Additional Outcome Specifier) to label them (e.g., "5 min post-ROSC", "30 min post-ROSC"). Flag shared animals in column 45 ("Same animals as [other row]"). This captures the temporal evolution of perfusion failure, which is important for the review. A transition must involve a change in the Perfusion Quality Category (column 41) --- specifically, a shift between a category indicating spatial flow presence (e.g., "No-reflow absent," "Hyperperfusion," "Hypoperfusion") and one indicating spatial flow failure (e.g., "No-reflow present (quantified)," "Qualitative flow absent"). Changes in flow *magnitude* that do not alter spatial patency (e.g., hyperperfusion at 10 minutes declining to hypoperfusion at 30 minutes, with flow still present in all regions at both times) do **not** qualify as a qualitative state change; extract only the earliest time point and document the trajectory in column 46.

**Consistency requirement:** Apply the qualitative state change exception uniformly across all experimental conditions within the same paper. If the same qualitative transition (e.g., no-reflow present → no-reflow absent) occurs in two different experimental conditions (different ischemia models, different animals in a case-series, etc.) and both transitions have supporting data, both warrant their own additional row. "Supporting data" means any evidence --- text description, figure data, or explicit group-level statement --- that the transition occurred and that the Perfusion Quality Category changed; it does not require quantitative precision at the transition time point. If the transition time point has only qualitative descriptions or figure-estimated values, create the row and populate the Perfusion Outcome column with whatever data is available (qualitative descriptions, figure estimates per §1.5), flagging limitations in the companion columns. Do not apply the exception selectively based on model eligibility concerns, sample size, data quality, data completeness, level of quantitative detail, or the need to rely on figure estimation rather than text-reported values --- those concerns are documented in the relevant Quality Notes and companion columns but do not affect row-creation decisions. This does not relax the requirement in §2.6 that the assessment method produce quantitative or semi-quantitative data --- the transition time point must still have an extractable numeric value (from text, table, or figure estimation per §1.5) in the Perfusion Outcome column, not a purely narrative description with no recoverable numbers. *Example 1:* If a paper uses both CSF compression and vascular occlusion models, each with cohorts at 5 and 60 min recirculation, and no-reflow resolves between those time points in both models, both models get two rows --- not one model with two rows and the other with one row plus a column 46 note. *Example 2:* In a case-series paper (§3.2A) where three individually reported animals each show a transition from no-reflow absent to trickle-flow present, all three animals get transition rows --- even if only one animal's transition is quantified with specific voxel percentages while the others are described qualitatively or require figure estimation. If you find yourself writing transition data into column 46 instead of creating a transition row that the consistency requirement demands, this is the extraction error the consistency requirement exists to prevent.

**Continuous monitoring data (e.g., ASL-MRI, continuous laser Doppler):** When perfusion is measured continuously rather than at discrete time points, the "earliest time point" is the first post-ROSC data point at which the perfusion signal has stabilized enough to be interpretable (i.e., is no longer dominated by the transient hemodynamic instability of the first 1--2 minutes). In practice, identify the first labeled time bin on the figure's x-axis after ROSC (often 2.5--5 min post-ROSC) and read the y-value at that x-coordinate, following the figure-estimation protocol in §1.5. Do not select a later time point where a drug effect peaks or where perfusion has evolved, even if the authors' statistical analysis focuses on that later window --- record those later findings in the Notes columns instead.

*Rationale:* Selecting a later time point where impairment has resolved would systematically underestimate initial perfusion failure and obscure evidence of reversible no-reflow. If the earliest stabilized time point shows hyperemia rather than hypoperfusion, extract it --- hyperemic readings demonstrate that perfusion is achievable and are valuable data for this review (see §0).

Document the recovery trajectory in column 46 (Notes - Long-Term Perfusion Improvement) rather than creating additional rows (except for the qualitative state change exception above).

### §3.2A Case-Series Papers with Individually Reported Animals

Some papers --- particularly early foundational studies --- present each animal as an individual experiment rather than as part of a pooled group with a shared protocol. When different animals within the same nominal experimental group were assessed at **different post-reperfusion time points**, create a **separate row for each distinct assessment time**, with each row containing only the animal(s) measured at that time. Do not pool animals measured at different times into a single row.

- If animal A was measured at 5 minutes and animal B at 30 minutes, these are two separate rows (each with n=1), not one row with semicolon-separated values.
- If animals A and B were both measured at 5 minutes, they share one row (n=2).
- Use column 4 (Additional Outcome Specifier) to label each row with the assessment time (e.g., "5 min post-reperfusion", "30 min post-reperfusion").
- Flag rows from the same study that share the same experimental conditions in column 45 ("Same experimental group as [other row], different assessment time").

**Distinguishing §3.2A from §3.2:** §3.2 applies in two situations: (1) when the **same set of animals** is measured serially (repeated measures), and (2) when **separate cohorts** under the same experimental condition are each assessed at a different time point (parallel-cohort design). In both cases → pick earliest, apply qualitative state change exception, note trajectory in column 46. §3.2A applies to a different scenario: **case-series papers** where different animals within a **single nominal group** happen to be measured at different times as individual experiments (independent observations → separate rows). If a case-series paper measures the same animal at multiple time points, apply §3.2 (pick earliest for that animal); if it measures different animals at different times, apply §3.2A (separate rows).

**Case-series interaction with qualitative state change:** In a case-series paper where each animal is measured serially, §3.2's qualitative state change exception applies *per animal*. If three animals in the same nominal group are each measured at multiple time points, and all three show a transition from no-reflow absent to no-reflow present, this produces up to **six** rows: one earliest-measurement row and one transition row per animal. The consistency requirement (above) applies across animals --- do not create the transition row for only the animal with the most detailed transition data while relegating the others to column 46. **The most common failure mode is recognizing that the consistency requirement applies but overriding it due to perceived disproportionality between row count and sample size, or because transition evidence for some animals is less precisely documented than for others. Neither is a valid reason to omit a transition row — data precision concerns belong in the companion columns, not in row-creation decisions.**

**Important:** This rule applies only to repeated measurements on the same group of animals. Groups that differ in ischemia duration, treatment, or assessment protocol are separate experimental groups, each requiring its own row.

#### §3.3 Temporal Scope

Only extract perfusion data from the early post-reperfusion period (generally within 30--60 minutes after restoration of circulation). Do not extract data points from hours or days post-reperfusion that primarily reflect secondary injury, delayed hypoperfusion, or long-term recovery. If the earliest available measurement is beyond 60 minutes, include it but note in the Quote column that no earlier measurement was available, and flag in Notes.

If a paper reports CBF serially over hours (e.g., at 15 min, 1 h, 3 h, 6 h), select the earliest representative time point.

#### §3.4 Do Not Split by Region or Metric

Do not create separate rows for different brain regions measured in the same group at the same time --- use the most comprehensive summary measure as the primary Perfusion Outcome and record region-specific data in Regional Breakdown (column 29).

Do not create separate rows for different metrics of the same phenomenon in the same group at the same time (e.g., both % no-reflow area and categorical pass/fail go in the same row across the appropriate columns).

#### §3.5 Different Treatments = Different Rows

Different treatment groups are separate rows even if ischemia duration is identical. This includes untreated or vehicle-control groups: if both groups underwent ischemia and had perfusion quality assessed, each gets its own row. **Do not relegate any group's perfusion data to the Notes column of another group's row** --- neither the control group's data into the treated row's Notes, nor the treated group's data into the control row's Notes. This applies even if the control data comes from a companion paper cited by the study being extracted.

**Small subgroups.** This rule applies regardless of subgroup size. When a paper reports individual-animal data for treatment subgroups with n=1 or n=2, each subgroup still gets its own row. Do not aggregate across different treatments to increase n. Small-n observations will be appropriately downweighted in the downstream meta-regression by inverse-variance weighting; the extraction should preserve the data as reported.

*Worked example:* A paper tests 10 min of ischemia in 4 untreated rabbits, 1 rabbit given heparin 4 mg/kg, 1 rabbit given heparin 10 mg/kg, and 1 rabbit given heparin 20 mg/kg --- all assessed at the same time point. This produces **4 rows** (one per treatment condition), not 1 row with treatment data in Notes. The n=1 heparin subgroups each get their own row.

#### §3.6 Sham and Control Groups

Do not create rows for groups that received no ischemia or only trivially brief ischemia intended as a control condition. Whether a group is a "control" depends on the study's own designation, not a fixed duration threshold --- a 3-min or 5-min ischemia group that is part of a genuine dose-response design (the study treats it as an experimental condition) should be extracted as its own row. Note sham/control perfusion values in the Notes column of the relevant experimental rows as baseline reference.

**Contrast with §3.5:** This rule excludes only groups with **no ischemia** (sham/controls). Groups that received ischemia but also received a drug treatment are **not** shams --- they are treatment groups and must have their own rows per §3.5, regardless of how small the subgroup is.

### §3.7 Multiple Assessment Modalities

**Prerequisite:** The multiple-modality rule applies only when each modality independently qualifies as perfusion quality data under §2.6. A method that provides supplementary hemodynamic or kinetic information but does not independently meet §2.6's qualification criteria should be documented in the Notes column of the qualifying modality's row, not extracted as a separate row. The tie-breaking rule in §3 does not override §2.6 eligibility --- a method that fails to qualify cannot generate a row regardless of the preference for more rows over fewer.

If a study uses multiple different methods to assess perfusion quality on the same cohort (e.g., carbon black perfusion and angiography, or gross assessment and CT), create a separate row for each method. "Independent" here means a different measurement technique, regardless of whether the measurements were performed on the same or different animals. Use column 4 (Additional Outcome Specifier) to label each row with the method name (e.g., "carbon black", "angiography"). Use the modality-specific sample size if reported; otherwise use the headline cohort N and note in the Quote column.

*Example:* A study subjects 10 animals to 10 min of cardiac arrest, then assesses perfusion quality using both carbon black perfusion and angiography. This produces two rows: one labeled "carbon black" and one labeled "angiography," both with n=10.

**Shared animals across rows.** When multiple rows from the same study share the same animals (whether from multiple modalities or other reasons), note this in column 45 (Notes): "Same animals as [other row identifier]." This flag enables the downstream multilevel meta-analysis to correctly model the within-study correlation rather than treating the rows as independent observations.

### §3.8 Post-Hoc Subgroups

If a paper subdivides a treatment group into subgroups based on an achieved perfusion-relevant variable (e.g., achieved reperfusion pressure, flow rate, temperature, functional recovery such as EEG return) and reports perfusion outcomes separately, treat each subgroup as a separate row --- provided the subgroups show categorically different perfusion outcomes. Do not split for non-perfusion variables (e.g., sex, weight) or only quantitatively modest differences. Label each subgroup in column 4 (Additional Outcome Specifier) with a descriptive label (e.g., "EEG recovery", "no EEG recovery", "high-pressure subgroup"). The column 4 label is the primary mechanism for distinguishing these rows; a separate Caveats flag is not required.

#### §3.9 Pharmacological Dose-Response and Parametric Manipulation Data

Some studies report CBF not as a standard observation at a defined post-reperfusion time point, but as part of a **dose-response or parametric manipulation experiment** --- e.g., CBF measured at a series of pharmacologically induced CPP levels, or CBF measured across a range of $PaCO_2$ values. The defining feature of such an experiment is that **a single independent variable is swept across multiple levels**, producing a curve rather than a single observation. These data characterize a dose-response relationship rather than the spontaneous post-ischemic perfusion state. **Do not create separate extraction rows for data points from such curves.** Extracting a value from a dose-response curve requires selecting an arbitrary operating point (e.g., "CBF at CPP ~100 torr"), which is a constructed observation, not a natural one.

Instead, summarize the key findings of the dose-response or parametric experiment in the Notes column (column 45) of the relevant experimental group's row. Include the range of the manipulated variable, the CBF response, and any notable findings (e.g., "CPP elevation from 64 to 105 torr had no significant effect on CBF during the post-ischemic low-flow state, Figure 1").

**What §3.9 does NOT cover:** A group of animals that received a defined single-protocol pharmacological treatment (e.g., a bolus of Drug X, or a sequence of Drug X followed by Drug Y) and had CBF measured at a defined post-treatment time point is a **treatment group under §3.5**, not a dose-response experiment under §3.9 --- even if the treatment was designed as a mechanistic probe, a specificity control, or a reversal experiment. The distinction is between sweeping a variable across multiple levels (§3.9) and applying a fixed treatment protocol to a cohort (§3.5). The motivation for the treatment (therapeutic intent vs. mechanistic inquiry) does not affect row-creation decisions; it is captured in the Caveats columns (§1.12).

**Interaction with §3.5 and §3.8:** If a subgroup of **the same animals** whose early data is already captured in the parent group's row underwent conditions that diverged from the main group *only after the primary observation time* (e.g., inadvertent hemodilution occurring after the standard serial measurements, or a vascular reactivity challenge such as acetylcholine or hypercapnia), and that subgroup's unique data exists only within a pharmacological challenge or manipulation experiment (whether a dose-response curve or a single-challenge test), the subgroup's data belongs in Notes rather than in a separate row. This clause applies only when the animals are literally a subset of another group's cohort, not merely when two independent cohorts happen to share the same pre-treatment experimental conditions. §3.5 applies to groups that differed in treatment during or before the ischemic and primary assessment period. It does not require a separate row for every animal subset whose conditions happened to diverge at a later experimental stage, particularly when no standard perfusion quality observation exists for the subgroup under its unique conditions.

**Consistency test:** Apply the same row-creation logic to all subgroups in the same experiment. If the main group's dose-response data (e.g., CBF autoregulation curve at normal hematocrit) is summarized in Notes rather than extracted as a separate row, then a subgroup's data from the same type of experiment (e.g., CBF autoregulation curve at reduced hematocrit) should also be summarized in Notes. Conversely, if you extract one single-protocol treatment group as its own row under §3.5, all other single-protocol treatment groups in the same study must also get their own rows --- you cannot apply §3.5 to one and §3.9 to another when both received a defined treatment and had CBF measured at a defined time point.

**Exception:** If a dose-response or parametric experiment is the *only* source of perfusion data for a group that underwent ischemia under distinct conditions from the start (e.g., a dedicated hemodilution treatment arm with no other perfusion assessment), §3.5 applies and a row should be created, selecting the operating point closest to normal physiological conditions. Note the dose-response context in the companion Quote column and in Notes.

### §4 Study Quality Table

One data row per paper. 17 columns (16 tabs per row). Do not include headers.

| # | Column | Instructions |
|---|---|---|
| 1 | **Study Authors** | First author last name + "et al." (or last name alone if sole author). |
| 2 | **Year** | Four-digit publication year. |
| 3 | **PMID** | PubMed ID, or "Not found." |
| 4 | **DOI** | Digital Object Identifier, or "Not found." |

| | | |
|---|---|---|
| 5 | **Ischemia Duration Clearly Defined?** | Yes / Yes -- Range / No / Unclear. "Yes" = a single exact duration is specified and experimentally controlled. "Yes -- Range" = the paper specifies a target duration but reports the achieved duration as a narrow range (e.g., "11--12 min"), indicating normal experimental variation around a controlled target. |
| 6 | **Ischemia Duration Quote** | Verbatim passage supporting the assessment. |
| 7 | **Sample Size Explicitly Reported?** | Yes / No / Unclear. "Yes" = the extractable experimental group size is explicitly stated either as a direct per-group n or as a narrative enumeration that unambiguously determines the group size (e.g., "three experiments with VF (Exp. 2-4)" for a paper that presents each experiment individually). "No" = only total cohort size is given and the size of the extractable group must be inferred indirectly. "Unclear" = the text is genuinely ambiguous about how many animals contributed to the reported outcome. **Granularity note:** This column assesses whether the paper reports sample sizes for its experimental groups as the paper defines them, not whether the per-row sample size for every extracted observation is determinable. If the paper states n=9 for a treatment arm but the Data Extraction creates multiple rows at different ischemia durations within that arm, answer "Yes" here (the paper reported the group size); the per-duration n issue is handled in the Data Extraction Sample Size column (col 7 of Data Extraction). |
| 8 | **Sample Size Quote** | Verbatim passage supporting the assessment. |

| | | |
|---|---|---|
| 9 | **Temperature During Ischemia Reported?** | Yes / No / Unclear. "Yes" if: (a) a specific temperature value is reported during ischemia (whether actively controlled or passively measured), OR (b) normothermic conditions were actively maintained throughout ischemia, OR (c) a specific temperature is reported immediately before ischemia onset and the ischemic period is brief enough (generally ≤15 min) that core temperature would not have changed meaningfully. If heating was discontinued during ischemia but a measured temperature is still reported (e.g., "body temperature declined to 35 ± 0.5°C"), score "Yes" --- the key criterion is whether the temperature is known, not whether it was controlled. "Unclear" if heating or temperature monitoring is mentioned but no specific temperature value during or immediately before ischemia is given (e.g., the paper mentions a rectal probe was placed but never reports what it read). "No" if no temperature information appears anywhere in the paper. |
| 10 | **Temperature Quote** | Verbatim passage reporting the temperature value or normothermic maintenance. "No supporting text" if assessment is "No." Each Quote column must contain ONLY the passage directly supporting the preceding assessment column. |
| 11 | **Perfusion Outcome Reproducibly Described?** | Yes / No / Unclear. "Yes" = methods detailed enough to replicate, including cases where the paper cites a prior publication for full methodological detail and provides enough information to identify the referenced protocol. If the referenced publication is genuinely needed to understand the methods (e.g., scoring criteria described only in the cited paper), pause and ask the user to provide that publication before proceeding. |
| 12 | **Perfusion Outcome Description Quote** | Verbatim passage supporting the assessment. |

| | | |
|---|---|---|
| 13 | **Blinding Used in Outcome Assessment?** | Yes / No / Unclear. If "Yes," specify scope in parentheses: "Yes (perfusion)", "Yes (histopathology)", or "Yes (perfusion and histopathology)". "Unclear" if blinding is not mentioned for any outcome --- absence of mention should be scored "Unclear," not "No." "No" only if the paper explicitly states blinding was not used. Automated or objective measurement methods (e.g., computed speckle contrast ratios, autoradiographic densitometry) do not require blinding and should not affect the assessment for other outcomes. |
| 14 | **Blinding Quote** | Verbatim passage supporting the assessment. |
| 15 | **Inclusion/Exclusion Criteria Stated?** | Yes / No / Unclear. "Yes" if the paper explicitly states prospective inclusion or exclusion criteria, explicitly reports post-hoc exclusions of specific analyzed animals with stated reasons (e.g., "one dog died and was excluded," "one animal was excluded because it experienced 15 minutes of ischemia instead of 13"), or describes protocol termination rules that removed animals from the analyzed cohort. "No" if no such analyzed-cohort rule or exclusion is stated. **Do not count mention of separate pilot studies, preliminary experiments, or unreported sham work as inclusion/exclusion criteria unless the paper explicitly says those animals were part of the analyzed cohort and then excluded.** "Unclear" only when the paper suggests but does not clearly state cohort-level exclusions. |
| 16 | **Inclusion/Exclusion Quote** | Verbatim passage supporting the assessment. |
| 17 | **Quality Notes** | Free text. If excluded, begin with: "EXCLUDED --- Model: [reason]", "EXCLUDED --- Outcome: [reason]", "EXCLUDED --- Design: [reason]", or "EXCLUDED --- Language: [reason]". For models requiring careful judgment (§2.5), include "Completeness confidence: High/Medium/Low" with justification. |

### §5 Data Extraction Table

One data row per observation (see §3). 47 columns (46 tabs per row). Do not include headers.

#### Identifier Columns (1–4, no companions)

| # | Column | Instructions |
|---|---|---|
| 1 | **Study Authors** | Same as Study Quality. |
| 2 | **Year** | Same as Study Quality. |
| 3 | **Link** | PubMed URL preferred, else DOI link, else "Not found." |
| 4 | **Additional Outcome Specifier** | Short label distinguishing this row from other rows for the same study (e.g., "untreated", "rt-PA treated", "high-pressure subgroup", "20 min ischemia", "gross assessment", "CT", "histology"). Leave blank if only one observation row. |

#### Paired Extraction Columns (5–34, alternating data + companion)

See §1.2 for the companion column convention. See §1.3 for data column formatting. See §1.4--§1.5 for data source and figure-estimation rules.

| # | Column | Instructions |
|---|---|---|
| 5 | **Species / Strain** | e.g., "rat (Wistar)", "cat", "human". |

| | | |
|---|---|---|
| 6 | **Species Quote** | |
| 7 | **Sample Size (n)** | Per-group n as a single number. "NR" if not reported; note total N if available. See §1.3 for handling uncertain or conflicting n values. When the paper reports a headline cohort size but some cases have missing perfusion data, use the headline size and note the effective n in the Quote column. |
| 8 | **Sample Size Quote** | |
| 9 | **Ischemia Duration (min)** | In minutes. Apply §1.1 (zero-flow boundary). If a range is reported for a single cohort (e.g., "20--25 min"), use the midpoint and note the range in the Quote column. For cardiac arrest with CPR, see §2.1. For asphyxia, see §2.2. When methods give a range but results state a single value, prefer the range (midpoint) and note the discrepancy. |
| 10 | **Ischemia Duration Quote** | |
| 11 | **Temperature During Ischemia** | Exact value if reported (e.g., "37°C rectal"). "Normothermic (assumed)" if not reported and no indication of hypothermia. "Hypothermic" with details if applicable. **Do not write "NR" for this column.** If no temperature information appears anywhere in the paper and there is no indication of hypothermia, the correct value is "Normothermic (assumed)" --- laboratory experiments on anesthetized animals default to normothermic conditions unless stated otherwise. |
| 12 | **Temperature Quote** | |

| | | |
|---|---|---|
| 13 | **Ischemia Model** | e.g., "cardiac arrest (KCl-induced)", "ascending aorta occlusion with IVC occlusion", "decapitation". For vascular occlusion models, specify all vessels occluded (see §2.5). |
| 14 | **Ischemia Model Quote** | |
| 15 | **Blood Flushed Prior?** | Yes / No / N/A / NR. "N/A" for in vivo blood reperfusion studies (e.g., cardiac arrest → resuscitation → blood circulates) where no separate perfusate is introduced. "No" for studies that introduce a perfusate or tracer but do not mention flushing --- blood flushing is a significant step that would be reported if performed. "NR" only if the protocol description is too vague to determine. |
| 16 | **Blood Flushed Quote** | |
| 17 | **Perfusate Type** | Be specific. Describe full sequence if multi-step. Include any pharmacological interventions that modify perfusion conditions (e.g., thrombolytics, heparin, hypertonic saline). |
| 18 | **Perfusate Quote** | |
| 19 | **Perfusion Pressure (mmHg)** | Report CPP (MABP − ICP) when both available; note component values in Quote. When ICP is not reported, report MABP and note in Quote that it represents MABP, not CPP. For ex vivo perfusion, report stated perfusion pressure directly. Apply §1.4 and §1.10 --- check hemodynamic tables, figures, **and protocol descriptions** (e.g., "MAP was maintained at >80 mmHg") before writing "NR." If reported in other units, convert to mmHg and note originals (§1.6). If pressure varied |

| | | |
|---|---|---|
| | | across animals, report the group mean or the pressure at which the primary comparison is made. |
| 20 | **Perfusion Pressure Quote** | |
| 21 | **Perfusion Quality Assessment Method** | e.g., "FITC-albumin fluorescence microscopy", "radioactive microsphere CBF". |
| 22 | **Assessment Method Quote** | |
| 23 | **Perfusion Outcome (Value)** | The most comprehensive summary measure (whole-brain or total-forebrain preferred). Report group mean ± variability when available. If only individual-animal data exists, list individual values separated by semicolons and note that no group summary was reported. For binned histograms, record the full distribution (e.g., "13/46 <20, 22/46 20--40, 11/46 >40 ml/100g/min"). Include units. Tag source per §1.4. For figure estimates, apply §1.5. **CBF values:** ALWAYS report in ml/100g/min; if the paper uses other units, convert (§1.6). **Spatial metrics:** report as % of assessed region with no-reflow or not perfused. **Capillary-level metrics:** keep as-is but note the different denominator. When individual-animal data is available in tables, report both individual values (semicolons) and the computed mean. |
| 24 | **Perfusion Outcome Quote** | |

| | | |
|---|---|---|
| 25 | **Variability (Type ± Value)** | Always explicitly state "SD" or "SEM". "NR" if none. "SD/SEM unspecified ± [value]" if the paper does not specify. Report variability in the same units as the Perfusion Outcome value (column 23); if a unit conversion was applied to column 23, apply the same conversion to the variability measure. Before writing "NR," check figure legends and methods for variability type declarations. If the paper reports only regional variabilities with no overall measure, write "NR" here and record regional variabilities in column 29. Apply §1.11: always quote the paper's exact descriptor in the companion Quote column. |
| 26 | **Variability Quote** | |

| | | |
|---|---|---|
| 27 | **% Brains Adequately Perfused** | Percentage of individual animals meeting a perfusion adequacy threshold. **Threshold selection:** Use the paper's own threshold when it defines one (e.g., "homogeneous reperfusion" = 0% no-reflow). When the paper does not define its own threshold, use a default of ≥90% spatial perfusion (i.e., ≤10% of assessed brain area with no-reflow or absent flow). State the threshold and its basis (e.g., "100% (5/5) using threshold: no-reflow ≤10% (default)", or "40% (2/5) using threshold: homogeneous reperfusion (paper-defined)"). Always compute from individual-animal data when available in tables rather than writing "NR." When a tracer study finds complete filling with no evidence of no-reflow in any animal, infer 100% and state the basis. Apply §1.9: when a spatially resolved method finds measurable flow in all assessed regions, infer 100% using "measurable CBF in all regions" as the threshold. For methods whose field of view does not cover the whole brain, write "NR" per the spatial resolution requirement in §1.9. Reserve "NR" for cases with no basis for estimation (e.g., only a single group mean CBF is reported with no individual or regional data). A group mean above zero does not by itself justify inferring 100%, because the mean could mask individual animals with zero flow. **Deriving from group-level statements:** When the paper provides explicit, unambiguous group-level statements about the consistency of a finding, use that statement to derive the percentage even if individual-animal data is not tabulated. Unambiguous quantifiers ("all," "every," "none," "no animals") can be converted directly --- e.g., "all animals showed homogeneous reperfusion" → 100% (n/n); "no-reflow was observed in every animal" → 0% (0/n); "all animals exhibited disturbed microcirculation" → 0% (0/n) using homogeneous reperfusion as threshold. **This rule applies symmetrically:** group-level statements establishing that all animals had impaired perfusion yield 0% just as reliably as statements establishing that all animals had intact perfusion yield 100%. Do not write "NR" when an unambiguous group-level statement provides a clear basis for derivation, even if individual-animal quantitative data is not tabulated. Ambiguous quantifiers ("most," "about half," "over a third") require the tag "(inferred from group-level statement, moderate confidence)." Always cite the verbatim statement in the Quote column. **Whole-brain inference safeguard.** Use 100% adequacy from group-level statements only when both of the following are true: (1) the method has whole-brain spatial relevance under §1.9, and (2) the paper makes an explicit group-level statement such as "no regional differences," "all regions showed flow," or equivalent. If either condition is missing, do not infer 100%; use "NR" unless another valid basis is available. |

| | | |
|---|---|---|
| 28 | **% Brains Quote** | |
| 29 | **Regional Breakdown** | Region-specific perfusion outcomes by *anatomical brain region* (cortex, hippocampus, basal ganglia, brainstem, cerebellum, thalamus, white matter, etc.). Format: "Region: value±variability; Region: value±variability". "N/A" if only whole-brain data. Do NOT use this column for flow-range bin distributions --- those belong in column 23 or Notes. **Unit preference:** When both absolute CBF values (ml/100g/min) and percent-of-control values are available for regional data, prefer absolute CBF in ml/100g/min. Use percent-of-control only when absolute values cannot be determined. State which unit is used. |
| 30 | **Regional Breakdown Source** | **Never skip this column.** "from text", "from Table N", "estimated from Fig. N (low confidence)", "estimated from Fig. N (very low confidence)", or "N/A". When column 29 is "N/A", this column must be "N/A". **Do not place explanatory prose in column 30.** Any explanation about why regional data are unavailable belongs in column 32, 45, or the relevant companion column, not here. |
| 31 | **Brain Region(s) Assessed** | Region in primary Perfusion Outcome (e.g., "total forebrain", "whole brain"). Also list additional regions from column 29. "Whole brain (assumed)" if unspecified. |
| 32 | **Brain Region Quote** | |

| | | |
|---|---|---|
| 33 | **Time from Ischemia Onset to Perfusion Quality Assessment (min)** | Total minutes from ischemia start to the moment perfusion quality is assessed --- not the time to the start of reperfusion. Apply §1.1 (zero-flow boundary). For tracer methods, the moment of assessment = tracer injection, not sacrifice. For continuous monitoring, the moment of assessment = the specific time point reported. For cardiac arrest with CPR, see §2.1 --- list all component intervals (VF, CPR, post-ROSC) in the Quote column. For figure time axes, see §2.4. For time-window averages, see §2.7. Write only a single number; show arithmetic in Quote column (§1.3). If intermediate reperfusion occurred before the final assessment, include the full timeline and describe the intermediate reperfusion in the companion Quote column. When the time is ambiguous, pick the midpoint and explain. |
| 34 | **Time to Perfusion Quote** | Quote passage(s) specifying the post-ischemia interval and moment of assessment. Show the arithmetic. |

### Derived/Interpretive Columns (35–44, alternating value + reasoning)

| # | Column | Instructions |
|---|---|---|
| 35 | **Blood vs. Non-Blood** | "Blood", "Non-Blood", or "Mixed". Refers to the medium circulating through the brain at the time of perfusion quality assessment. Blood = brain perfused with blood when assessment occurs (includes in vivo microsphere studies --- microspheres are tracers within blood). Non-Blood = brain perfused with non-blood solution, or tracer perfused through blood-free vasculature. Mixed = combination at time of assessment. |
| 36 | **Blood vs. Non-Blood Reasoning** | Explain classification. |

| | | |
|---|---|---|
| 37 | **Caveats Category** | Apply §1.12. Select all that apply, semicolon-separated: "None beyond standard resuscitation" / "Pre-treatment (experimental intervention)" / "Pre-treatment (standard experimental anticoagulation)" / "Intervention to improve perfusion quality" / "Intervention that worsens perfusion" / "Post-resuscitation vasopressors" / "Hypothermic ischemia" / "Other (specify)". |
| 38 | **Caveats Details** | Describe specific interventions, including drug names, doses, and timing. "N/A" if Caveats Category is "None beyond standard resuscitation". |
| 39 | **Time from Perfusion Onset to Perfusion Quality Measurement (min)** | Minutes between reperfusion onset (ROSC, clamp release, perfusate flow start) and the moment of assessment. Apply §1.1 (zero-flow boundary). "NR" if not determinable. "0" if assessment was at the moment of reperfusion. For post-mortem tracer perfusion after in vivo reperfusion, "perfusion onset" = when blood flow was restored, not when tracer was injected later. For time-window averages, see §2.7. |
| 40 | **Time from Perfusion Onset to Measurement Quote** | Show arithmetic and source passage. |
| 41 | **Perfusion Quality Category** | Apply §1.8 (no-reflow priority rule). One of: "No-reflow present (quantified)" / "No-reflow absent" / "Hypoperfusion (quantified CBF)" / "Hyperperfusion (quantified CBF)" / "Qualitative flow present" / "Qualitative flow absent" / "Mixed regional heterogeneity". Write only the category label. **Low-pressure non-filling:** When assessment uses a tracer at sub-physiological pressure (e.g., <50 mmHg), quantified non-filling is still "No-reflow present (quantified)." Note in column 42 that non-filling may reflect inadequate pressure rather than (or in addition to) vascular obstruction. |

| | | |
|---|---|---|
| 42 | **Perfusion Quality Category Reasoning** | Brief explanation. **Required sub-qualifier when PQ Category is "No-reflow present (quantified)":** State "Sub-qualifier: fixed" (consistent with permanent capillary obstruction --- e.g., post-mortem tracer filling defects), "Sub-qualifier: dynamic/reversible" (consistent with vasospasm or transient dysfunction --- e.g., low-flow regions that migrate or resolve), or "Sub-qualifier: persistence unknown" (study design does not permit distinction --- e.g., single time-point assessment). |
| 43 | **Converted % Brain Perfused** | Apply §1.9. Integer 0--100 (never a 0--1 decimal). "Cannot convert" if not feasible. Do NOT convert from methods whose field of view does not cover the whole brain (see §1.9). Do NOT invent volumetric estimates from qualitative spatial descriptions ("small" or "large" zones) unless a figure allows visual estimation of affected fraction. |
| 44 | **Conversion Reasoning** | Show conversion method or explain why not feasible. |

#### Terminal Notes Columns (45–47, no companions)

| # | Column | Instructions |
|---|---|---|
| 45 | **Notes** | Edge cases, ambiguities, figure-only data, sham/control baseline values, excluded groups (e.g., RCP groups per §2.5), additional context. |
| 46 | **Notes - Long-Term Perfusion Improvement** | Actively look for and document any evidence that perfusion impairment resolved or improved over time during reperfusion. Check ALL serial time points in the paper, including later time points from figures and tables not extracted as the primary observation. Include specific time points and values with supporting quotes. E.g.: "No-reflow areas at 5 min recirculation had disappeared by 60 min |

| | | |
|---|---|---|
| | | (Table 2), replaced by delayed hypoperfusion." Do not leave blank without first checking all serial data for recovery evidence. |
| 47 | **Notes - Regional Differences in Perfusion Impairment** | Systematic regional differences in perfusion quality (e.g., certain brain regions consistently perfusing better or worse, spatial patterns of no-reflow distribution). Summarize with supporting quotes. Leave blank if not applicable. |
