## Supplementary material for "How long is the brain perfusable after global ischemia? A systematic review": DataS7_Frequency-of-adequate-perfusion.html


### Frequency of adequate perfusion

**Perfusate:** 
● Pure blood 
● Carbon black 
● Mixed blood 
● Other non-blood

**Shape:** 
● n reported 
▲ n not reported

**Size:** 
larger points = larger n per arm

Hover over points for study details. Click a point to open the source paper.
